# Mantis: high-throughput 4D imaging and analysis of the molecular and physical architecture of cells

**DOI:** 10.1101/2023.12.19.572435

**Authors:** Ivan E. Ivanov, Eduardo Hirata-Miyasaki, Talon Chandler, Rasmi Cheloor-Kovilakam, Ziwen Liu, Soorya Pradeep, Chad Liu, Madhura Bhave, Sudip Khadka, Carolina Arias, Manuel D. Leonetti, Bo Huang, Shalin B. Mehta

**Author notes:** These authors contributed equally to this work (Ivan E. Ivanov, Eduardo Hirata-Miyasaki, Talon Chandler, Rasmi Cheloor-Kovilakam).

## Abstract

High-throughput dynamic imaging of cells and organelles is essential for understanding complex cellular responses. We report Mantis, a high-throughput 4D microscope that integrates two complementary, gentle, live-cell imaging technologies: remote-refocus label-free microscopy and oblique light-sheet fluorescence microscopy. Additionally, we report shrimPy, an open-source software for high-throughput imaging, deconvolution, and single-cell phenotyping of 4D data. Using Mantis and shrimPy, we achieved high-content correlative imaging of molecular dynamics and the physical architecture of 20 cell lines every 15 minutes over 7.5 hours. This platform also facilitated detailed measurements of the impacts of viral infection on the architecture of host cells and host proteins. The Mantis platform can enable high-throughput profiling of intracellular dynamics, long-term imaging and analysis of cellular responses to perturbations, and live-cell optical screens to dissect gene regulatory networks.

**Significance Statement:** Understanding the dynamics and interactions of cellular components is crucial for biological research and drug discovery. Current dynamic fluorescence microscopy methods can only image a few fluorescent labels, providing a limited view of these complex processes. We developed Mantis, a high-throughput 3D microscope that maps interactions among components of dynamic cell systems. Mantis combines light-sheet fluorescence imaging of multiple fluorophores with quantitative label-free microscopy and is complemented by shrimPy, our open-source software for high-throughput data acquisition and high-performance analysis. Mantis enabled simultaneous 3D time-lapse imaging of 20 cell lines and quantitative analysis of responses to perturbations like viral infection at single-cell resolution. This approach can accelerate the analysis of cellular dynamics and image-based drug discovery.

## Introduction

Several open problems in cell biology and drug discovery require high-throughput methods to measure, predict, and model the dynamic interactions among cells, organelles, and proteins. Multiplexed imaging of cellular morphology and machine learning are increasingly used to predict the effects of genetic and chemical perturbations (1–3) from changes in the cellular architecture. High-throughput live cell imaging, combined with deep learning, has enabled systematic analysis of the dynamic mechanisms that underpin healthy (4–6) and diseased (7–9) states of cells.

Correlative imaging of the 3D architecture of cells and constituent components over time (4D imaging) can accelerate discovery and dissection of cellular processes. However, 4D imaging of more than three organelles with multispectral fluorescence microscopy (10) remains challenging, because engineering cells with multiple fluorescent proteins is labor intensive and the wide emission spectra of fluorescent proteins limits the number of labels that can be imaged simultaneously. Correlative label-free and fluorescence microscopy is a viable strategy to mitigate the longstanding multiplexing bottleneck in 4D microscopy, because label-free imaging captures many cellular structures at once. Several cellular landmarks, e.g., nuclei, cell membrane, and nucleoli, scatter sufficient visible light and can be consistently visualized with label-free imaging. Several organelles, e.g., cytoskeleton, endoplasmic reticulum, and Golgi, as well as individual proteins need to be labeled with fluorophores for consistent visualization as they do not scatter visible light measurably in live cells. Correlative label-free and fluorescence imaging systems have recently been reported (4, 11–14) for 3D imaging of organelles, cells, and tissues.

Building on these advancements, we have developed an automated microscope, named "Mantis", that synergizes light-sheet and label-free microscopy in multiwell plates. Light-sheet and label-free microscopy are both gentle on live samples as they do not require high light doses that can cause phototoxicity. We multiplex oblique light-sheet fluorescence microscopy (15–19) with remote refocus (20, 21) quantitative label-free imaging with phase and polarization (QLIPP) using distinct wavelengths. Mantis enables 4D imaging of 3 or more fluorescent (i.e. extrinsic) labels and 3 physical properties of phase, retardance, and orientation as described in (4) at high speed and in parallel. The large field of view of the microscope and the combination of two imaging modalities provides rich phenotypic data on the dynamics of specific molecules in the context of the cell architecture. The name “Mantis” is inspired by the high-dimensional vision and the quick reflexes of the mantis shrimp.

Automated, robust, and configurable acquisition software is essential for 4D imaging, especially in high-throughput format. High-throughput correlative microscopes can produce 10s of terabytes of data per day. Parallelized, configurable, and reproducible analysis is essential to leverage the statistical patterns captured with these large datasets. We integrate high-throughput acquisition and high-performance computing, streamlining the process of imaging and profiling at high resolution up to a million cells in a single experiment. The acquisition and analysis engine is implemented in python and is available as open source repository (shrimPy) on GitHub (22). shrimPy streamlines calibration of the optical path, deconvolution of the specimen properties from the acquired intensities, and segmentation of single nuclei and cells. Instead of experimentally staining nuclei and cell membranes, we rely on virtual staining of quantitative phase images (4, 23). Virtual staining of nuclei and membrane frees up fluorescence channels for imaging of proteins and other organelles, improves the imaging throughput, and counteracts loss of fluorescence due to photobleaching or stochastic labeling. shrimPy analysis engine uses our recent robust models for joint virtual staining of nuclei and membrane (23).

We demonstrate the performance and applications of Mantis by reporting 3D tracking of organelles and parallel imaging of 20 cell lines expressing distinct fluorescently-tagged intracellular reporters-in the context of cellular landmarks that are imaged without labels. We illustrate joint virtual staining of nuclei and membrane, and accurate single cell segmentation in multiple cell lines. Gentle imaging with Mantis enables analysis of the temporal dynamics of the cell morphology and the localization of key host proteins in response to viral infection. Used together, these tools enable high-content analysis of the dynamics of intracellular reporters in the context of the global morphology of the cell.

## Results

### Multichannel 4D imaging of cell dynamics

The Mantis microscope is a synergistic combination of oblique light-sheet fluorescence microscopy and remote refocus quantitative label-free imaging with phase and polarization (QLIPP) as illustrated in Figure 1a and detailed in <u>Methods</u>. The integration of two gentle imaging modalities through wavelength multiplexing enables simultaneous imaging of both the physical and molecular compositions of dynamic biological samples at high-throughput in multiwell plates. The label-free measurements enable gentle imaging of many organelles in terms of the distribution of their density and biomolecular orientation. Quantitative label-free imaging provides key landmarks for subsequent segmentation and preserves the fluorescence spectrum for labeled proteins and organelles.

**Figure 1.**
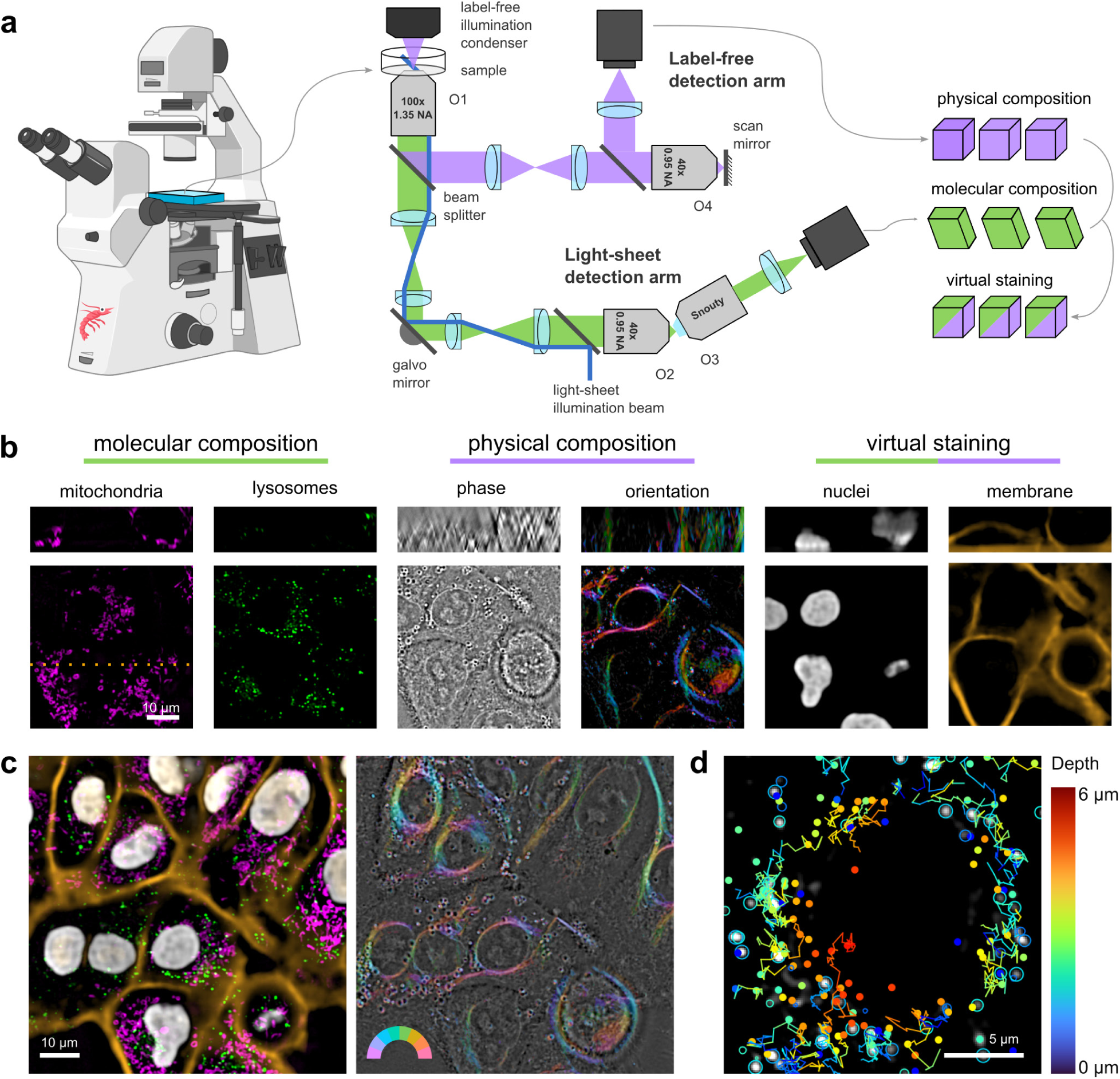
Overview of the Mantis microscope. (a) Schematic of the light path and data acquisition pipeline. Mantis enables high-throughput measurements of the physical and molecular composition of dynamic biological samples in multiwell plates by combining remote-refocus label-free microscopy and oblique light-sheet fluorescence microscopy. Landmark organelles such as nuclei and membranes are virtually stained from label-free images. (b) Images of A549 cells in different channels. Fluorescent and virtually stained organelles can be captured in the context of the overall cell architecture. Left to right: labeled mitochondria (TOMM20-GFP) and lysosomes (LysoTracker), label-free measurements of phase and molecular orientation, and nuclei and plasma membrane virtually stained from the phase channel. Top row shows XZ orthogonal projections of data presented in the bottom row at the location indicated by the orange dotted line. (c) Correlative imaging of physical and molecular composition enables 4D imaging of multiple organelles in parallel. Left: overlay of mitochondria (magenta), lysosomes (green), nuclei (white), and membrane (orange); right: overlay of phase (grayscale) and orientation (color). (d) Time-lapse imaging enables tracking of organelles. Line segments show the movement of lysosomes in 3D (depth encoded in color) over time. Circles show in-focus spots, dots show out-of-focus spots.

Long-term imaging of intracellular dynamics and image-based screens in a multiwell plate format are key applications of the Mantis platform. We chose oblique light-sheet fluorescence microscopy as it does not impose additional sample mounting requirements, reduces phototoxicity by only illuminating the plane in the sample that is being imaged, and enables fast volumetric imaging by scanning a galvo mirror (16, 18). The principle of remote refocus microscopy (15, 20, 21) underpins this design-the oblique light-sheet and detection perpendicular to the light-sheet are implemented in the remote volume. To enable fast correlative label-free imaging, Mantis speeds up the axial scanning in label-free phase and polarization microscopy (4) by an order of magnitude using remote-refocus architecture.

The primary objective of the microscope is a 100x 1.35NA silicone immersion lens (O1, Figure 1a) and is shared by the label-free and fluorescence arms. The remote refocus path in both arms is implemented using 40x 0.95NA air objective (O2 and O4) and suitable tube lenses to achieve magnification of 1.4x, equal to the ratio of refractive indices of silicone oil and air, as dictated by the principles of remote refocus (20, 21). The remote volume in the fluorescence arm is imaged with a “Snouty” objective (O3), which is perpendicular to the oblique light-sheet (17). The remote-refocus objective (O4) in the label-free arm is reused to magnify the remote volume after reflection from a scan mirror. A detailed optical schematic and CAD models of the microscope are available in Figure 1-Supplement 1, Figure 1-Supplement 2, and Figure 1-Supplement 3. In this specific configuration, the Mantis microscope enables volumetric imaging of over ∼15 μm depth without moving the primary objective or the sample because axial scanning is carried out entirely in the remote-refocus arms. The current light-path can be configured to image up to 60 μm deep volumes by changing the angle between light-sheet and coverslip, and the corresponding angle between imaging axes of O2 and O3.

Building a microscope that achieves high resolution, high light efficiency, and minimal polarization distortions required the development of the following optical alignment procedures and optical modules: A) a method for attaching a glass coverslip to the O2 and O4 objectives to compensate for spherical aberrations (Figure 1-Supplement 4); B) a polarized-light analyzer cube that is insensitive to birefringence of the dichroic beam splitter (Figure 1-Supplement 5); C) a polarization-based label-free remote refocus path that maximizes light throughput (Figure 1-Supplement 6); D) a procedure for calibrating the 3D point spread function (PSF) of the light-sheet path using beads distributed in 3D agarose gel (Figure 1-Supplemental Note 1). These procedures correct for majority of the optical aberrations, enable diagnosis of the alignment, and provide calibration data needed to deconvolve the residual aberrations. These procedures play a critical role in enabling high throughput dynamic imaging with diffraction-limited resolution over multiple days, and facilitate re-alignment as needed, typically once a month.

We illustrate the channels of information that can be acquired with Mantis using A549 cells (Figure 1b and Figure 1-Supplemental Movie 1). We visualized mitochondria and lysosomes by tagging TOMM20 with GFP and staining the cells with LysoTracker Deep Red. The phase channel of the same field of view reports the spatial distribution of biomolecular density and visualizes the nuclei, nucleoli, plasma membrane, and other dense organelles. The orientation channel reports the orientation of angular order (4) among biomolecules and visualizes plasma membrane and perinuclear fibers that appear similar to vimentin filaments previously reported in this cell line at high confluence (24–26). Lastly, we used virtual staining to predict nuclei and plasma membrane from the phase image (<u>Methods-Virtual staining and segmentation</u>). Virtual staining of these landmark molecular markers can enable segmentation and single-cell analysis using existing segmentation models built for fluorescence images (23). All channels of information can be acquired in 3D-in Figure 1b we show orthogonal slices through the cell body with distinct morphological structures evident throughout the volume.

We overlay fluorescence and label-free images in Figure 1c and Figure 1-Supplemental Movie 2. We show the entire 150 x 150 µm field of view of the Mantis microscope in all channels of information in Figure 1-Supplement 7. We acquired these data at a rate of 2 volumes per minute, which enabled us to do 3D tracking of lysosomes (Figure 1d and Figure 1-Supplemental Movie 3).

### High-throughput acquisition and analysis

Robust high-content imaging of multiple proteins in the context of the global cell morphology required the development of a dedicated acquisition and analysis engine (Figure 2) that we named shrimPy (Smart High-throughput Robust Imaging & Measurement in Python) (22).

**Figure 2.**
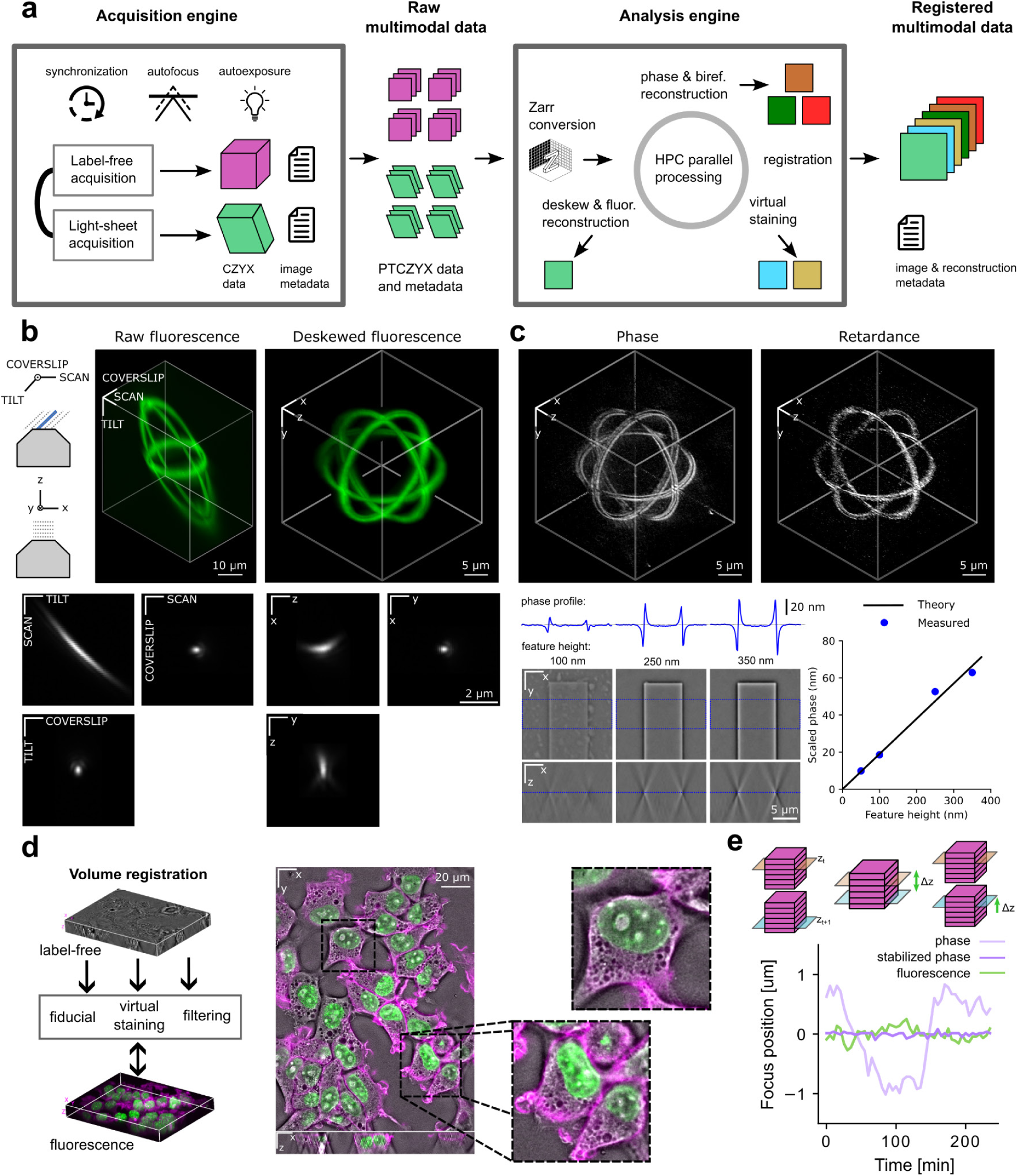
Automation for high-throughput data acquisition and image processing. (a) The shrimPy acquisition engine synchronizes data collection from the two arms of the microscope, acquiring images over time and positions, and is responsible for autofocus and autoexposure. Raw label-free and light-sheet PTCZYX datasets (P: position; T: time; C: channel; Z, Y, and X: three spatial dimensions) are converted to Zarr format for input to shrimPy analysis engine. Light-sheet data are deconvolved and deskewed, label-free data undergo phase and birefringence reconstruction, and cellular landmarks are predicted using virtual staining of phase images. Processing of the data is accelerated by a high-performance cluster (HPC). Metadata is generated and traced through each step of the pipeline. Data originating from the two arms of the microscope are registered to generate correlative multimodal datasets. (b) Deskewing of light-sheet fluorescence volumes. Top left: an oblique light sheet illuminates the sample and defines the raw fluorescence sampling coordinates; after deskewing we show data in Cartesian coordinates aligned with the objective optical axis. Middle and right: raw and deskewed 3D maximum intensity projections of Argolight sphere target. Bottom: raw and deskewed slices through a representative bead point-spread function. (c) Reconstruction of label-free volumes. Top: maximum intensity projections of phase and retardance reconstructions of the Argolight sphere target. Bottom: quantitative phase reconstructions of 100, 250, and 350 nm high glass features; a linear relationship between feature height and reconstructed phase is seen. (d) Registration of label-free and fluorescence volumes. Data from different modalities can be registered in 3D by incorporating fiducial markers visible in all channels, by virtual staining of structures also visible in the fluorescence channels, or by applying filtering operations (e.g. Sobel filter) to both datasets to maximize their mutual information. (e) Axial stabilization of the label-free and light-sheet volumes. The light-sheet arm is stabilized by a hardware autofocus feedback loop. The label-free arm is stabilized computationally post acquisition by subtracting the focus drift estimated from all positions at a given time point (see Methods).

The shrimPy acquisition engine orchestrates parallel data acquisition from the label-free and light-sheet arms and carries out smart microscopy tasks such as autofocus and autoexposure. The two arms of the microscope run on two instances of Micro-Manager (27, 28), which we control using the Pycro-Manager python bridge (29). The shrimPy acquisition engine coordinates their synchronous operation using hardware triggering (Figure 2-Supplement 1 and <u>Methods-Microscope automation</u>).

The acquired multidimensional raw datasets are processed by the shrimPy analysis engine to generate registered multimodal data that can be used for phenotyping (Figure 2a). Raw data are first converted into the OME-Zarr format (30), which enables efficient parallel processing of multiple timepoints and positions. As described below, discrete data volumes then undergo deskewing of fluorescence channels, reconstruction of phase and orientation, registration, and virtual staining. Parallel processing is accelerated using our high-performance computing cluster (Figure 2-Supplement 2).

Light-sheet volumes are acquired in a skewed frame of reference due to the oblique illumination and detection (Figure 2b and Figure 2-Supplement 3) and need to be deskewed for visualization in the frame of reference of the imaging chamber and for downstream analysis alongside the label-free data. Figure 2b shows the transformation of a fluorescent sphere target (Argolight) between skewed (here SCAN-TILT-COVERSLIP) coordinates and deskewed (X-Y-Z) coordinates. After deskewing, the target assumes a spherical shape as expected (Figure 2-Supplemental Movie 1). We also show images of sub-resolution fluorescent beads in both coordinate systems. These data demonstrate optical resolution of 290 nm in SCAN, 260 nm in COVERSLIP, and 290 nm in TILT; or 290 nm in X, 260 nm in Y, and 680 nm in Z coordinates (see Figure 2-Supplement 4 and Figure 2-Supplemental Movie 2). We improve our resolution and contrast further by applying a bead-based deconvolution routine (see Figure 2-Supplement 5), where we average beads to measure the microscope PSF, apply a Tikhonov-regularized least-squares deconvolution, and then deskew the results.

Phase (optical path length), retardance (difference of optical path length between the two symmetry axes of the structure), and orientation (orientation of the axis along which the biomolecular density is higher) are reconstructed from raw brightfield or polarized light images as described earlier (4). The microscope was calibrated for polarized light imaging and the acquired data reconstructed using the recOrder plugin (31) for napari. Figure 2c shows phase and retardance reconstruction of the same Argolight sphere. The target has a different refractive index than its surrounding medium and is weakly birefringent, showing signal in our label-free detection channels. Figure 2-Supplement 6 further shows projections of the sphere in the three principal planes in each of the three contrast modes. To our knowledge, our work reports the quantitative imaging with phase and polarization using remote-refocus acquisition for the first time. In Figure 2c glass features of increasing height were imaged to demonstrate that the measured phase increases linearly with the height of features, i.e. with increasing optical path length, as expected. We quantify the transverse (XY) spatial resolution for phase imaging to <400 nm (Figure 2-Supplement 7) — note that as in fluorescence imaging, the ability to resolve objects depends on both their density and the detection signal-to-noise ratio, here governed by the difference in refractive index between the objects and their surrounding medium.

Registration between the label-free and fluorescence volumes acquired on the two arms of the microscope is critical for correlative data analysis. Although volume registration can be pre-calibrated using targets such as Argolight or fluorescent beads embedded in agarose gel, the registration between the two arms can drift over days. However, the registration remains stable during the time required to acquire all positions. To enable robust registration from the acquired volumes we have developed following strategies (Figure 2d): A) We include fiducial markers such as large fluorescent microspheres that produce strong signal in both the label-free and fluorescence channels as one position in the multi-position acquisition. We used this approach to register data shown in Figure 1. B) In some experiments, fiducial markers are not needed, because molecular markers present in the fluorescence channel can be virtually stained (4, 23) from phase volumes. In these cases, nuclei or plasma membranes are predicted from phase volumes and the 3D similarity transformation matrix is computed by maximizing mutual information between the experimental and virtual stain (Figure 2-Supplement 9). C) In experiments where the fluorescence channel encodes membranous organelles, image processing filters (e.g. edge detection) applied to both modalities suffice to achieve sufficient mutual information for registration.

Enabling long-term experiments requires compensating mechanical drift between the image planes of the four objectives of the Mantis microscope (Figure 2e). The primary objective (O1) can be kept in focus using the built-in Nikon Perfect Focus System. The drift between the O2 and O3 objectives in the light-sheet remote refocus arm (see Figure 1a) leads to misalignment of the illumination and detection planes, which leads to substantial degradation of image quality. To maintain the alignment between the O2 and O3 objectives, an image-based autofocus method (Figure 2-Supplement 8) is implemented in the shrimPy acquisition engine (Figure 2e, green trace). Axial drift between the remote-refocusing objective (O4) and the scan mirror was also observed in the label-free arm. However, the drift in the label-free arm leads to imaging of different depths of the sample rather than a loss of image contrast. Therefore, the drift in the label-free arm is stabilized as a post-processing step (Figure 2e, compare dark and light purple lines). A hardware image-based autofocus method may also be developed in this arm of the microscope to reduce the need for imaging outside of the axial region of interest.

### Live cell phenotyping

The shrimPy acquisition and analysis engine enables high-throughput correlative measurements of physical and molecular architecture in a multiwell plate format over several hours. Figure 3 shows phase and fluorescence images of 20 cell lines from the OpenCell library (32) expressing a unique endogenously-tagged protein and a CAAX–mScarlet reporter of the plasma membrane. Volumetric imaging of fluorescently tagged proteins (Figure 3-Supplemental Movie 1 and Figure 3-Supplemental Movie 2) spanning multiple organelles (nuclei, nucleoli, Golgi, ER, endosomes, lysosomes, cytoplasm, stress granules) illustrates the ability of Mantis and shrimPy to map the localization of diverse targets with single-cell resolution.

**Figure 3.**
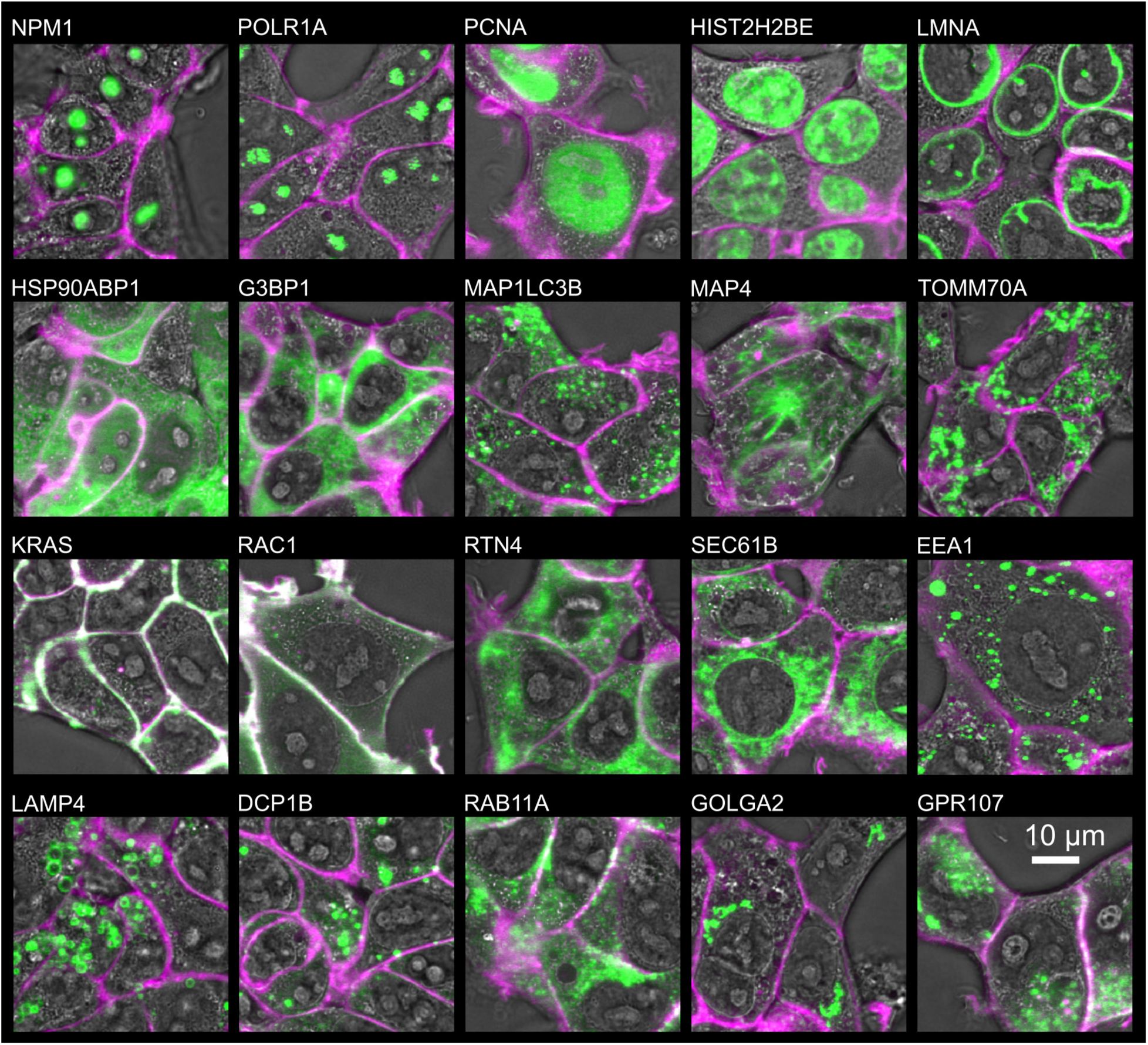
High-throughput imaging using the Mantis microscope. Images of 20 OpenCell targets with overlaid phase (grayscale), CAAX-mScarlet labeled membrane (magenta), and split-mNeonGreen2 tagged molecular marker (green) channels. Fluorescent channels are shown as a maximum intensity projection over a 1.1 μm z-section of the sample.

Live cell phenotyping typically requires segmentation of nuclei and cytoplasm in order to parse heterogeneity of cellular responses to the perturbations. We leverage virtual staining (23, 33) to enable image based phenotyping with single-cell resolution (Figure 4). Recently, generalist models for segmenting nuclei (34–36) and membrane (36, 37) of diverse cell types have been developed. They perform well for fluorescence images but require substantial human annotation to adapt to label-free datasets (38). Optimizing these models to segment label-free images would require onerous human annotation effort, especially for 3D segmentation. Virtual staining bypasses the need for human annotations, instead using molecular labels for annotation. We leveraged our recently published robust virtual staining model VSCyto3D (23), and CellPose (36) for single-cell segmentation (<u>Methods-Virtual staining and segmentation</u>).

**Figure 4.**
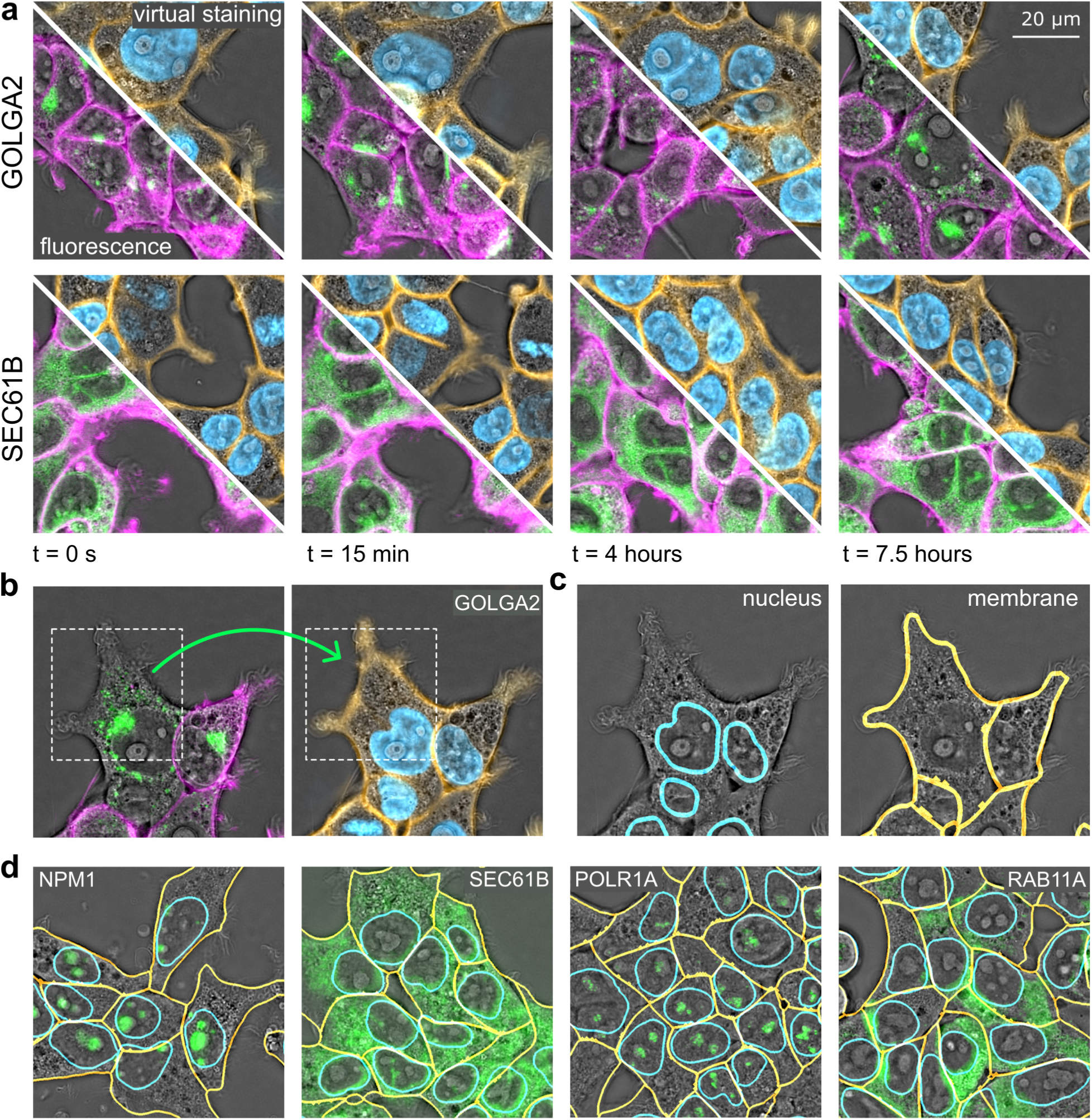
Virtual staining and instance segmentation of nuclei and membrane. (a) Comparison of membrane stained with CAAX-mScarlet (magenta, bottom-left half of FOVs) and virtually stained membrane (orange, top-right half of FOVs) in SEC61B and GOLGA2 cell lines (green) indicates reliable virtual staining. Virtually stained nuclei (blue) provide an additional channel of information. (b) Virtual staining rescues missing membrane labels. White bounding box and the green arrow show cells in the GOLGA2 line that do not express the CAAX-mScarlet label, which are rescued by virtual staining of the membrane. (c) Segmentation of virtually stained nuclei and membrane facilitates single-cell analysis of the localization and expression of fluorescently tagged proteins. Contours of the nuclear and membrane masks are shown in cyan and yellow, respectively. (d) Examples of virtual staining-based segmentation of nuclei and membrane of the cell lines in the OpenCell library.

VSCyto3D model led to reliable prediction of membrane and nuclei as illustrated in Figure 4a and Figure 4-Supplemental Movie 1. Note that VSCyto3D model was trained with widefield deconvolution dataset, and generalized to HEK293T volumes acquired with the label-free arm of the Mantis microscope without any fine-tuning. In some cell lines, a fraction of cells had lost their fluorescent membrane marker. Experimental and virtually stained images of one such FOV are shown in Figure 4b. The virtual staining model, which learns correlations between the two imaging modalities, rescued the missing label. By the same argument, virtual staining models cannot learn uncorrelated noise. Therefore, virtually stained fluorescence images are intrinsically denoised. Thus, virtual staining relaxes the experimental constraints on cell line engineering and live cell imaging.

Segmentation of virtually stained nuclei and membranes is shown in Figure 4c and Figure 4d for five cell lines in which proteins with diverse localizations are labeled: GOLGA2 (localized to Golgi), NPM1 (localized to nucleoli), SEC61B (localized to ER), POLR1A (RNA Polymerase, localized to nucleoli), and RAB11A (localized to recycling endosomes). To confirm the accuracy of segmentation of virtually stained nuclei and membrane, we imaged the HIST2H2BE cell line expressing fluorescently-labeled histone and plasma membrane markers. As shown in Figure 4-Supplement 1, the segmentations obtained from experimental label and virtual staining show high correspondence as measured by common metrics used to evaluate instance segmentation (<u>Methods-Virtual staining and segmentation</u>).

The modular pipeline consisting of reconstruction, virtual staining, and segmentation, described above enables the analysis of intracellular dynamics across multiple cell lines and perturbations with single-cell resolution.

### Analysis of viral infection dynamics

The speed and gentleness of the Mantis microscope enables molecular and morphological profiling of cells over long time periods, even when subjected to perturbations that induce cell stress. To assess this capability, we imaged cells from the OpenCell library that were infected with the common cold human coronavirus hCoV-OC43 (OC43). We chose OC43 as a model virus due to its disease relevance and experimental ease of use (39–41), and we chose to image from 20 to 50 hours post infection (hpi) because cells undergo large molecular changes during those times as measured by comprehensive organellar immunoprecipitation-mass spectrometry (42). In-depth follow up of these molecular changes, however, remains challenging due to phototoxicity induced by live-cell imaging using confocal microscopy.

Mantis enables long-term imaging with minimal phototoxicity, and thus allows close monitoring of protein dynamics in response to infection and identification of image-based viral infection sensors. Given that coronaviruses use the endoplasmic reticulum (ER) to replicate, which leads to eventual activation of the unfolded protein response pathway (40, 41), we monitored protein folding chaperons. We discovered substantial and consistent change in the localization of protein folding chaperon HSP90AB1 during infection. We virtually stained cell membrane and nuclei, and segmented cells using CellPose (see Figure 4 and <u>Methods</u>). This approach allows image based profiling of changes in protein localization, cell morphology, and cell density with single cell resolution.

Figure 5 shows the impact of OC43 infection on the cell morphology and the localization of HSP90AB1. The phase images show that infected HSP90AB1 cells are more condensed and have sharper variations in dry mass (Figure 5a, 40 hpi), relative to the mock condition. This observation is further supported by decrease in cell number and increase in the phase interquartile range (IQR), which demonstrates condensation of cell mass that is typically found in distressed or dying cells (Figure 5b and Figure 5-Supplemental Movie 1). In the fluorescence channel, we observed increased protein condensation and formation of puncta, which we quantified by measuring the skewness of the fluorescence intensity distribution in the cell cytoplasm and the number of puncta per cell (<u>Methods-Calculation of phenotypic features</u>). To our knowledge, condensation of HSP90AB1 in infected cells has not been directly observed before. The phase IQR and fluorescence skewness measurements are positively correlated, and allow clustring of infected and uninfected cells through straightforward gaussian mixture model using single-cell measurements from a short time-window (Figure 5-Supplement 1). The clustering algorithm enables classification of infection state of single cells from the phase IQR and fluorescence skewness features across the infection time course (Figure 5c and Figure 5-Supplemental Movie 2). These data indicate that quantitative phase images and their features may enable label-free readout of the cell’s state of infection, enabling high-throughput quantitative analysis of how organelles respond to infection.

**Figure 5.**
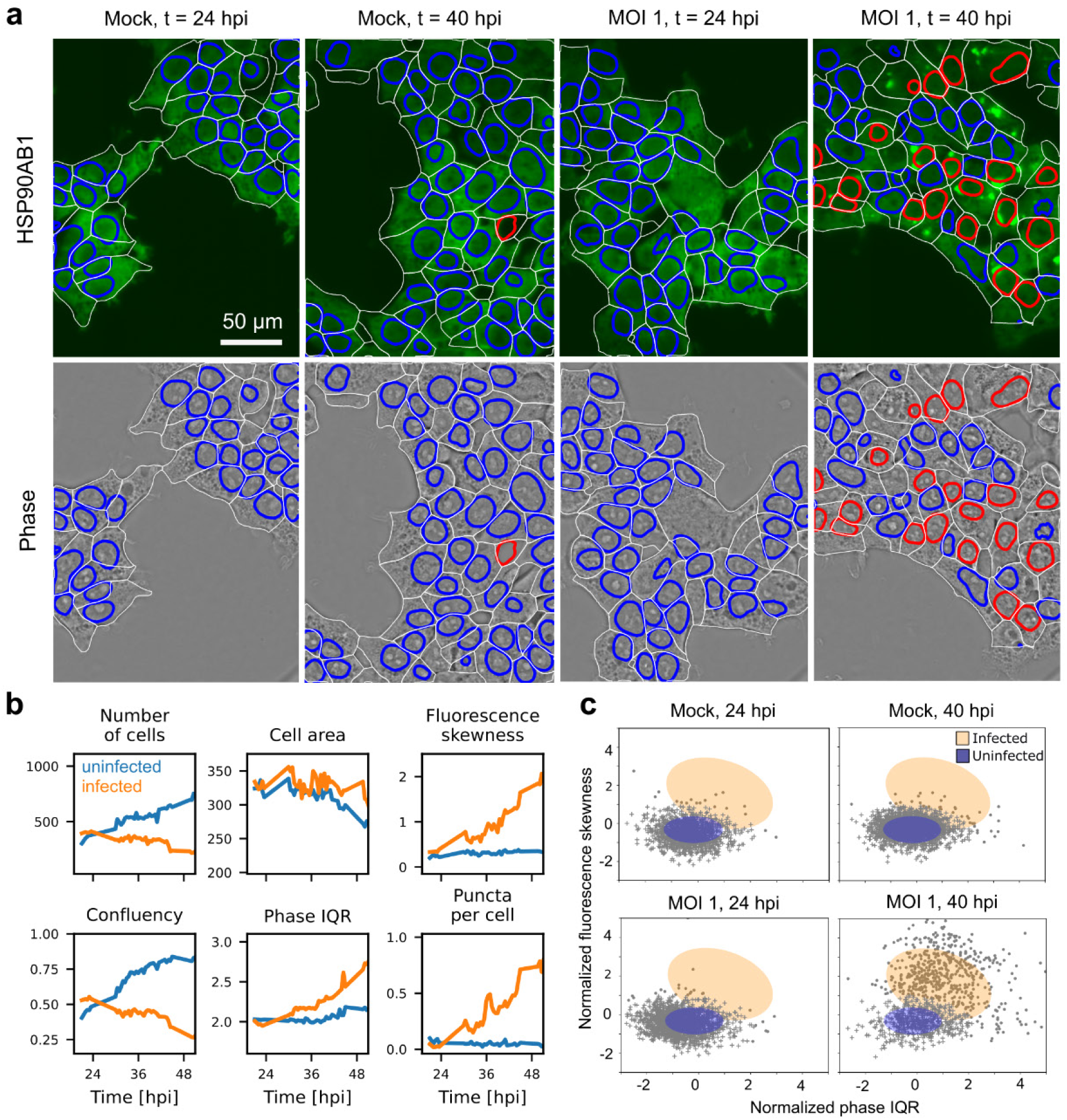
Single-cell phenotyping of OC43 infection. (a) Infected state prediction in images of phase (bottom half of FOVs) and fluorescence (top half of FOVs, maximum intensity projection over 4.1 µm z-section) in the HSP90AB1 control cells (top left, mock infection) and cells infected with OC43 ß-coronavirus (top right, MOI 1) at 24 and 40 hours post infection (hpi). Uninfected cells marked by blue nuclear boundaries and infected cells by red nuclear boundaries. (b) Measurements of phenotypic markers showing changes over the time course of infection. Membrane and nuclei of single cells were segmented after virtual staining as in Figure 4. The membrane mask was used to calculate the number of cells, cell confluency, cell area (in µm^2^), the interquartile range (IQR) in phase (in milliradians), and the skewness in fluorescence intensity. We further segmented and counted the number of large HSP90AB1 aggregates from fluorescence images (see Methods) and show the average number of puncta per cell over time. (c) Classification of infected and uninfected population of cells in mock and infected conditions using Gaussian mixture model. Uninfected cell population is covered by the blue Gaussian ellipse and marked by ‘+’ marker and infected population by orange ellipse and ‘o’ markers.

## Discussion

We described an automated imaging and analysis platform for 4D high-throughput imaging of molecular components and physical properties of cells and organelles. The Mantis imaging system consists of wavelength-multiplexed oblique light-sheet fluorescence microscopy and remote-refocus quantitative label-free microscopy. The shrimPy acquisition and analysis engine enables high-throughput imaging and profiling of intracellular dynamics. We demonstrate the spatial resolution and contrast with test targets, the temporal resolution with dynamic imaging of A549 cells, and high-throughput imaging capabilities by following 20 cell lines from the OpenCell library over time. Correlative data obtained with Mantis and shrimPy can be leveraged for simultaneous virtual staining of nuclei and membrane. Virtual staining and instance segmentation of nuclei and membrane enabled single-cell analysis and relaxed the experimental constraints on cell line engineering and live cell imaging. This platform enabled long-term imaging and profiling of changes in protein localization and cell morphology under stress induced by viral infection. Instrument design specifications as well as software for automation, image reconstruction, and virtual staining are shared via open-source repositories. Next, we discuss how we elected to balance the design tradeoffs, promising directions for subsequent technological developments, and key applications of the platform.

We currently acquire fluorescence channels sequentially. The oblique plane illumination geometry maps the oblique slice through the sample volume to a short strip of the camera sensor. By adding an image splitter before the camera that maps fluorescence channels to separate areas of the sensor, two or three emission channels can be imaged in parallel, improving the throughput and temporal resolution. Furthermore, the light-sheet and label-free arms acquire data at the same volumetric rate at present. However, the photon budget in the label-free arm is sufficient to operate it 2-5x faster than the light-sheet arm with few enhancements in automation. By using fast-switching liquid crystal devices (43) to modulate the polarization state of illumination light and improved synchronization sequences, we can drive the components of the label-free arm faster than the light-sheet arm. Lastly, we opted for a silicone immersion objective to minimize spherical aberrations when imaging deep into biological specimens. We can integrate recent design improvements (44) in remote-refocus microscopy for aberration-free imaging with silicon or air immersion with small adjustments to the light path.

The tight integration between acquisition and image analysis creates opportunities of improving the quality and utility of image data with computational microscopy, without increasing the complexity of the hardware. We have reported online alignment of two objectives in the detection path of the light-sheet arm that was essential for keeping the acquisition plane aligned with the illuminated plane of the sample. We can extend the approach to online label-free selection of fields of view with sufficient cell density and long-term tracking of collective cell dynamics.

Although high-throughput analysis is essential for iterative pursuit of biological questions, it remains significantly more challenging than high-throughput imaging. We have made algorithmic and engineering design choices in an attempt to make high-throughput analysis as tractable as high-throughput imaging. We opted for the OME-Zarr format throughout our image reconstruction and analysis pipeline to enable parallel processing of the data and efficient visualization with N-dimensional visualization tools, such as napari (45). All the steps in our analysis workflow are written such that they can be scaled up on a high-performance computing cluster. We developed focus estimation algorithms, PSF calibration algorithms, 3D registration algorithms, and 3D deconvolution algorithms. We integrated 3D virtual staining and 2D instance segmentation of cells as the first step in single cell analysis. 3D segmentation and tracking of biological structures remains an exciting open opportunity. We reported long-term imaging, single cell segmentation, and image-based profiling of infected cells. These image-based phenotypes can now be leveraged to identify label-free markers of infection.

The state-of-the-art imaging and analysis throughput of the Mantis platform creates new opportunities for systematic mapping and analysis of how diseases reprogram cells. For example, we are pursuing systematic mapping of how viral infections or genetic perturbations reprogram organelles by leveraging a subset of fluorescently tagged organelles from the OpenCell library. These dynamic measurements, parsed with suitable machine learning models that leverage temporal dynamics, can clarify the mechanisms of disease and lead to informative markers for drug screens.

The throughput of the Mantis platform can be adapted for several other applications. For example, we and others have shown that label-free imaging of antibody-stained (4) and H&E-stained (4, 14) tissue sections can improve the reliability and speed of imaging the tissue architecture. The high-speed correlative imaging capabilities of Mantis platform can accelerate digital pathology studies. Another exciting application is correlative analysis of the dynamics of molecules, organelles, and cells with sub-second temporal resolution. Correlative imaging of signaling dynamics, cell division, and cell differentiation is valuable for addressing open questions in developmental and cell biology. This adaptation can enable study of outstanding questions in how the biological function at higher spatial scales emerges from interactions at finer scales.

## Conclusion

In summary, we report measurement of molecular dynamics in the context of biomolecular density and orientation using Mantis, a 4D correlative imaging system that integrates label-free and light-sheet microscopy. In combination with an integrated acquisition and analysis engine (shrimPy), Mantis enables scalable analysis of image data with subcellular resolution. The Mantis platform enables longitudinal 3D imaging of multiple cell lines expressing multiple molecular markers in the context of the cellular morphology. We demonstrate the ability to track 3D dynamics of organelles in live cells with high temporal resolution. We also demonstrate long-term imaging and quantitative profiling of changes in protein localization and cell morphology of infected cells. We anticipate that our approach will enable systematic mapping and analysis of biological processes that govern the health and disease states of cells in diverse biological systems.

## Methods

### Microscope layout

The Mantis microscope is built around a Nikon Ti2 microscope body. Label-free imaging is performed at 450 nm, and longer wavelengths are used for fluorescence excitation and detection. The two detection arms of the microscope are separated using a dichroic beamsplitter positioned under the objective. The light-sheet remote-refocus arm is built following the design in Sapoznik et al. (18). The remote-refocus arm for label-free microscopy was designed by combining concepts described in Guo et al.(4) and Botcherby et al.(20). A detailed optical schematic is provided in Figure 1-Supplement 1, and the mechanical layout of the optical paths is given in Figure 1-Supplement 2 and Figure 2-Supplement 3. For birefringence imaging, the sample was illuminated with circularly polarized light using a universal polarizer (46) positioned near the back-focal plane of the microscope condenser; when only phase imaging was performed, the sample was illuminated with unpolarized light. For inspection of the sample before imaging we used the widefield epi-illumination and side detection port built into the microscope body. The primary (O1) objective was kept at a fixed position and the desired focal plane was selected by moving the sample axially using a piezo plate on the microscope stage or using custom spacers and sample holders when larger translation was needed.

### Microscope automation

Data acquisition is accomplished using custom python software (see Figure 2a) (47). The shrimPy acquisition engine coordinated parallel acquisition from the two remote-refocus arms of the microscope, collecting data over time and positions, and performed smart microscopy tasks such as autofocus and autoexposure. Each of the remote-refocus arms of the microscope was controlled by an independent instance of Micro-Manager (27, 28) and Pycro-Manager (29), collecting data over channels and z-slices. The two parallel acquisitions were synchronized using trigger pulses generated by a National Instruments DAQ card (NI cDAQ-9178). The exposure pulses of each camera were routed to a dedicated TriggerScope (AVR Optics) which changed the state of hardware associated with that acquisition (see Figure 2-Supplement 1).

### Sample preparation and imaging

The Argolight target used in this study was model Argo-SIM. The quantitative phase target was purchased from Benchmark Technologies and immersed in water for imaging. To measure the microscope point spread function 0.1 μm FluoSpheres beads (Thermo Fisher Scientific, F8803) embedded in 0.5% low melting point agarose (Sigma-Aldrich A2576) dissolved in 50% (w/w) glycerol in water were excited at 488 nm and imaged using a 525/45 nm bandpass filter.

CRISPR/Cas9 was used to generate endogenously tagged A549 TOMM20-GFP cell line (Figure 1). Briefly, wild type A549 cells (ATCC CCL-185) were electroporated with a mixture of *S. pyogenes* Cas9 protein (Macrolab), sgRNA targeting the protein of interest (PMID: 35271311 (32), GAGCTTGGCTGAAGATGATG, IDT) and a double-stranded homology-directed repair donor (42) using Amaxa 96-well shuttle nucleofector (Lonza) according to manufacturer’s protocol. Cells were allowed to recover in media with 1 µM nedisertib (M3814; Selleckchem # S8586) for 2 days. Electroporated cells were then expanded and sorted using SONY FACS to sort GFP positive cells.

HEK293T cell lines from the OpenCell (32) library were labeled with H2B-mIFP and CAAX-mScarlet using standard lentivirus transduction protocols.

Engineered HEK293T and A549 cells were cultured in 37°C and 5% CO2 and maintained between 20% and 90% confluency. Cells were routinely grown in DMEM with GlutaMAX (Thermo Fisher Scientific, 10566024), 10% fetal bovine serum (Omega Scientific, FB-11), and 100 U/mL penicillin/streptomycin (Thermo Fisher Scientific, 15140163). For imaging, HEK293T cells were seeded on 96-well glass-bottom plates (Greiner Bio One, 655891) coated with 50 µg/mL fibronectin (Corning, 356008, following manufacturer’s protocol); A549 cells were seeded on 12-well glass-bottom plates (Cellvis, P12-1.5H-N).

Prior to imaging, the microscope PSF was benchmarked as described in Figure 1-Supplemental Note 1 and the O1 correction collar was adjusted to minimize the PSF full width at half-maximum. Optionally, we collected datasets of immobilized beads and fluorescein solution for fluorescence deconvolution and flatfield correction.

Cells were imaged in DMEM media without phenol red (Thermo Fisher Scientific, 21063029) supplemented with 10% fetal bovine serum, 5 mM L-glutamine (Thermo Fisher Scientific, 25-030-081), and 100 U/mL penicillin/streptomycin. Cells were maintained at 37 °C and 5% CO2 using an OkoLab stage-top environmental chamber (H301-K-FRAME). Prior to imaging, the cell culture media was supplemented with ProLong Live antifade reagent (Thermo Fisher Scientific P36975) at 1:66 ratio. A549 cells were stained with 100 nM LysoTracker Deep Red (Thermo Fisher Scientific L12492). We used polydimethylsiloxane (PDMS) oil with 12500 cSt viscosity (MicroLubrol) as immersion medium for the primary (O1) objective as it provided better coating of the sample chambers. A549 cells were imaged using elliptically polarized light with swing of 0.05 waves as described in (4); HEK293T cells were imaged using unpolarized illumination to reconstruct the quantitative phase.

### Virus propagation and infection

OC43 (ATCC, VR-1558) were propagated in Huh7.5.1 cells at 34°C. To determine viral titers, 800,000 BHK-T7 cells per well were seeded in 6-well plates for 24h at 34°C. Virus stocks were then 10-fold serially diluted and applied onto cells for 2 hours at 34°C. Media was then replaced with DMEM containing 1.2% Avicel RC-591. Infected cells were incubated for 6 days at 34°C, then fixed with 4% formaldehyde, and stained with crystal violet for plaque counting. All experiments were performed in a biosafety level 2 laboratory. Aliquots of OC43 stored in-80°C were used to infect HEK293T cells with a target multiplicity of infection (MOI) of 1. Day prior to infection, 8000 HEK293T cells were seeded on 96-well glass-bottom plates (Greiner Bio One, 655891) coated with 50 µg/mL fibronectin (Corning, 356008, following manufacturer’s protocol) in DMEM with GlutaMAX (Thermo Fisher Scientific, 10566024), 10% fetal bovine serum (Omega Scientific, FB-11), and 100 U/mL penicillin/streptomycin (Thermo Fisher Scientific, 15140163). On day of infection, media was replenished with the same growth media with or without OC43 (viruses were thawed on ice). For MOI calculation, cells were assumed to have tripled. Final volume in a 96 well plate well was 100 µL.

### Image processing

Raw fluorescence data were deskewed by applying an affine transformation with trilinear interpolation resampling (Figure 2-Supplement 3). The affine transformation was parameterized by two quantities: the ratio of the object-space pixel width to the object-space scan step and the light sheet tilt angle. These parameters were estimated in three different ways: (1) for Figure 1 and the beads in Figure 2(b) we used a bead sample imaged before and after known stage motions along the X and Z axes, (2) for the Argolight target in Figure 2(b) we chose the affine transformation that restored the spherical shape of the target (Figure 2-Supplement 6), and (3) for the remaining figures we used the transformation estimated from the Argolight target followed by an axial rescaling to account for the mismatched RI = 1.52 index of refraction. Imaging into an index-mismatched target like the Argolight sphere leads to depth-dependent axial scaling (48), so we empirically measured our axial rescaling factor to be ∼1.3. Following deskewing, every three axial slices were averaged, improving SNR without losing resolution (Figure 2-Supplement 3).

Optionally, we applied a bead-based deconvolution to our fluorescence data to improve resolution and contrast (Figure 2-Supplement 5). As a calibration step, we acquired a volume of beads, found bead patches, and averaged these patches into a single PSF. We then used the averaged PSF to apply a Tikhonov-regularized least squares deconvolution to our data before deskewing. We show deconvolved and deskewed results in Figure 1, Figure 1-Supplemental Movie 1, Figure 1-Supplemental Movie 2, Figure 1-Supplemental Movie 3, and Figure 2-Supplement 5; elsewhere we show results without deconvolution. We find our simple bead-averaging method for estimated PSFs performs well on the datasets we tested, but we expect performance to degrade in very dim and sparse samples where inverse modeling techniques are more suitable (49).

Label-free data were reconstructed using recOrder (31). We generated a physics-informed model of the label-free image formation process, providing linear mappings between object properties (phase, retardance, and orientation) and the measured image intensities (4). Retardance and orientation were estimated by applying a pseudoinverse algorithm to five intensity measurements acquired under different polarized illuminations. For A549 data shown in Figure 1, retardance and orientation were further processed using a 3×3×5 median filter. Phase was estimated using a Tikhonov-regularized least-squares inverse algorithm applied to brightfield volumes.

Label-free and deskewed fluorescence volumes were registered by choosing the 3D similarity transformation (translation, rotation, and scaling) that maximized the mutual information between virtually stained nuclei or membrane and the corresponding fluorescent target using the Advanced Normalization Tools (ANTS) library (50). The optimization algorithm was initialized with a coarse manually chosen transformation (Figure 2-Supplement 9). The resulting transformation was then applied to all label-free volumes in the dataset.

We used a stabilization procedure to eliminate undesired motion in the registered image volumes over time. Label-free volumes were stabilized axially by finding in-focus slices by maximizing the transverse midband power for each time point and position, averaging over positions, then applying the averaged shift to each volume (Figure 2e).

### Virtual staining and segmentation

The virtual staining of nuclei and cell membrane from phase was done using the VSCyto3D model (23) fine-tuned using the mantis A549 cells. The fine-tuning dataset was composed of 100 FOVs of (9µm,147µm,124µm) ZYX volumes or (40,985,835) pixels. The model checkpoint with the lowest validation loss value was used for prediction and evaluation. Prediction from phase images of A549 and HEK293T cells was performed with the VisCy pipeline (23). Phase images were normalized to zero median and unit interquartile range prior to inference. The volumes were predicted in 5 z-slice sliding windows, and then blended by computing the mean of all windows. We share the models we have used for virtual staining and instance segmentation of nuclei and membrane via our GitHub repository, VisCy (33).

The nuclei and membranes of HEK293T cells were segmented using CellPose, a generalist algorithm for cellular segmentation. The 2D segmentation of the membrane was done using the built-in *‘cyto2*’ CellPose model without modifications by providing both the membrane and the nucleus as input channels. The 2D segmentation of the nucleus was done by extending the built-in Cellpose *‘nuclei*’ model. Our model, ‘CP_20220902_NuclFL’, uses additional manual annotations on fluorescence images of HEK293T and A549 cells with Hoecht stain (51).

We confirmed the performance of virtual staining models for the segmentation tasks by comparing the instance segmentations of the experimentally labeled nuclei and cell membrane and the virtually stained nuclei and cell membrane (Figure 4-Supplement 1). We used HIST2H2BE-mNeonGreen cell line in which the cell membrane was labeled with CAAX-mScarlet. We computed DICE coefficient per FOV to assess the degree of overlap between segmented nuclei and segmented cell membrane. We also computed average precision of detecting instances of nuclei or cells for each FOV. In some FOVs, the average precision was found to be low, which turned out to be the FOVs in which some cells were missing the HIST2H2BE-mNeonGreen or CAAX-mScarlet label. Subsequently, these models were applied to segment virtually stained nuclei and membrane and perform single cell analysis (Figure 4 and Figure 5).

### Calculation of phenotypic features

Phenotypic features shown in Figure 5 were calculated based on the cell membrane mask and image quantities in the phase or fluorescence channels. The number of cells was calculated as the total count of membrane masks in the 9 fields of view acquired for each condition. The confluency was calculated as the fraction of image size covered by membrane masks. Cell area was calculated on a per-cell basis using the membrane mask and averaged across all cells. Fluorescence skewness and phase interquartile range (IQR) were similarly calculated on a per-cell basis from the respective channels and averaged across all cells. Puncta in the fluorescence images were detected by intensity thresholding, erosion of the binary mask to exclude very small regions, and applying a determinant of Hessian blob detector using scikit-image.

### Infection state classification

Cell infection state as shown in Figure 5 was predicted using the Gaussian mixture model. Single cell HSP90AB1 skew and phase IQR were computed from 4.1 um max projection of HSP90AB channel and a single slice of phase image. Single cell information was achieved combining the cell segmentation information for each FOV of imaging, and each condition of imaging.

Gaussian mixture model was trained on normalized HSP90AB1 skew and normalized phase IQR data from infected condition from time 41.5 to 43.5 hpi. Gaussian fitting on infected and uninfected cell populations in mock and infected conditions were performed at the training window (figure 5 supplement image). The infection class was predicted for every cell on images from the training window to visually validate the results.

The model was applied to time indices 24 hpi and 40 hpi (Figure 5) on mock and infected conditions. Gaussian fitting of the infected and uninfected populations were also performed on both conditions at both time points. Single cells were classified as infected or infected based on the classification.

### Tracking

Tracking of lysosomes shown in Figure 1 was performed using TrackMate 7 in Fiji (52). We used a Laplacian of Gaussians spot detector and the Simple LAP tracker to generate lysosomes trajectories.

## Data availability

Datasets reported in this manuscript will be posted to Bioimage Archive after the peer review is completed. Virtual staining and instance segmentation of nuclei and membrane are available via GitHub (33).

## Acknowledgments

We thank Shivanshi Vaid, Grace Yun, and Vincent Turon-Lagot, CZ Biohub San Francisco, for preparing cell samples. We thank Rachel Banks for contributing the segmentation models used for segmenting experimentally and virtually stained nuclei and membranes. We thank Sandra Schmid for careful reading of the manuscript and critical feedback.

## Supplemental Information

**Figure 1-Supplement 1.**
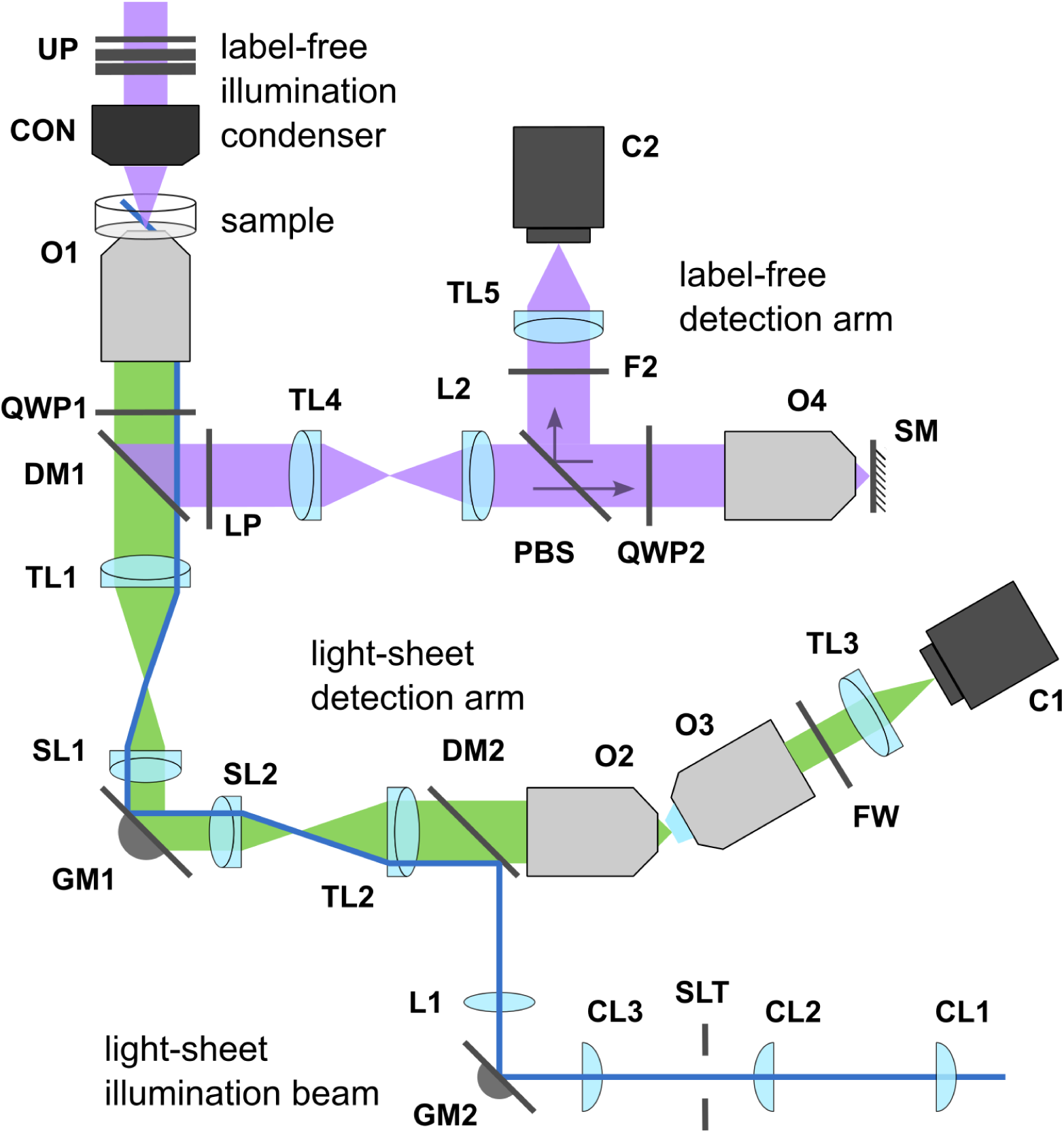
Simplified optical schematic. Light from the sample is collected using a 100x 1.35 NA silicone oil immersion objective (O1: Nikon Instruments Inc. MRD73950, f = 2 mm). Two remote-refocus paths are built to allow for oblique detection of fluorescence (green beam) and fast axial scanning in label-free imaging (purple beam). The two paths are separated by wavelength using a long-pass dichroic beamsplitter (DM1: Chroma Technology Corp. T475lpxr-UF2) positioned under O1. The primary image plane in the light-sheet detention arm is formed by the tube lens built into the Nikon Ti2 microscope body (TL1). The image plane is then relayed using two scan lenses (SL1: Thorlabs Inc. CLS-S, f = 70 mm and SL2: Thorlabs Inc. LSM03-VIS, f = 39 mm) and a galvo mirror (GM1: Thorlabs Inc. GVS201). A remote volume with 1.4x magnification is formed using a tube lens (TL2: Thorlabs Inc. TTL200, f = 200 mm) and a 40x 0.95 NA objective (O2: Nikon Instruments Inc. MRD70470, f = 5 mm). An oblique plane in the volume is then imaged using the “Snouty” objective (O3: Applied Scientific Instrumentation AMS-AGY v1, f = 5 mm and TL3: Thorlabs Inc. TTL200, f = 200 mm) on a fast back-illuminated sCMOS camera (C1: Teledyne Photometrics Prime BSI Express). Fluorophore emission filters (Semrock FF01-525/45-25-STR, FF01-600/37-25-STR, FF01-698/70-25-STR) are loaded on a filter wheel (FW: Sutter Instrument Company LB10-NWE) placed after O3. The light-sheet excitation beam (blue) is coupled into the optical path using a dichroic beamsplitter (DM2: Chroma Technology Corp. ZT405/488/561/640rpcv2-UF1) positioned behind O2 and a lens (L1: Thorlabs Inc. AC254-075-A). The excitation beam was formed by coupling the output of three laser sources (Vortran Stradus 488 nm, 561 nm, and 639 nm) into an optical fiber (not shown). The fiber output was then collimated (Thorlabs Inc. RC04FC-P01), expanded (using lenses AC254-050-A and AC254-150-A), and shaped into a linear profile using an achromatic cylindrical lens (CL1: Thorlabs Inc. ACY254-100-A). The beam was relayed using two additional cylindrical lenses (CL2: Thorlabs Inc. ACY254-050-A and CL3: Thorlabs Inc. ACY254-100-A) and a slit (SLT: Thorlabs Inc. VA100C) between them was used to adjust the numerical aperture of the light sheet illumination. For label-free imaging, the sample was illuminated with 450 nm light from a LED light source (Thorlabs Inc. M455L4 with a Semrock FF02-447/60 cleanup filter and Edmund Optics 54504 diffuser) and a Nikon 0.52 NA condenser (CON). The light was polarized with a controllable universal polarizer (UP), composed of a linear polarizer (Thorlabs Inc. WP25L-UB) and two variable retarders (Meadowlark Optics Inc.), oriented as described before (46). Polarized light transmitted by the sample was analyzed by a quarter-wave plate (QWP1: Thorlabs Inc. WPQ10M-445) and a linear polarizer (LP: Thorlabs Inc. WP25M-UB) positioned, respectively, before and after DM1 (see Figure 1-Supplement 5). The primary image plane in the label-free detection arm was formed with a tube lens (TL4: Thorlabs Inc. TTL200-A, f = 200 mm) and relayed with a composite lens (L2, built from Thorlabs Inc. lenses ACT508-500-A and ACT508-750-A as described in (17), f = 357 mm) and a 40x 0.95 NA objective (O4: Nikon Instruments Inc. MRD70470, f = 5 mm) to form a remote volume with 1.4x magnification. A scan mirror (SM: Edmund Optics 34-386 mounted on a Mad City Labs Nano-OP65 piezo stage) was placed in the remote volume to enable fast axial imaging, reflecting light back towards O4. A polarizing beam splitter (PBS: MOXTEK, Inc. FBF04C-UF) and a quarter wave plate (QWP2: Thorlabs Inc. AQWP10M-580) were placed behind O4 to maximize the throughput of the optical path and reflect light towards the detector (see Figure 1-Supplement 6). An image was formed using tube lens TL5 (Thorlabs Inc. TTL165-A, f = 165 mm) and recorded on a fast machine vision camera (C2: Teledyne Flir LLC Oryx ORX-10GS-51S5M-C). A bandpass filter (F2: Semrock FF01-445/20) before TL5 was used to reject stray light. For clarity this schematic omits steering mirrors and positioning stages which can be seen in Figure 1-Supplement 1 and Figure 1-Supplement 2. We glued No. 1.5 glass coverslips (Thorlabs Inc, CG15NH1) to objectives O2 and O4 as described in Figure 1-Supplement 4.

**Figure 1-Supplement 2.**
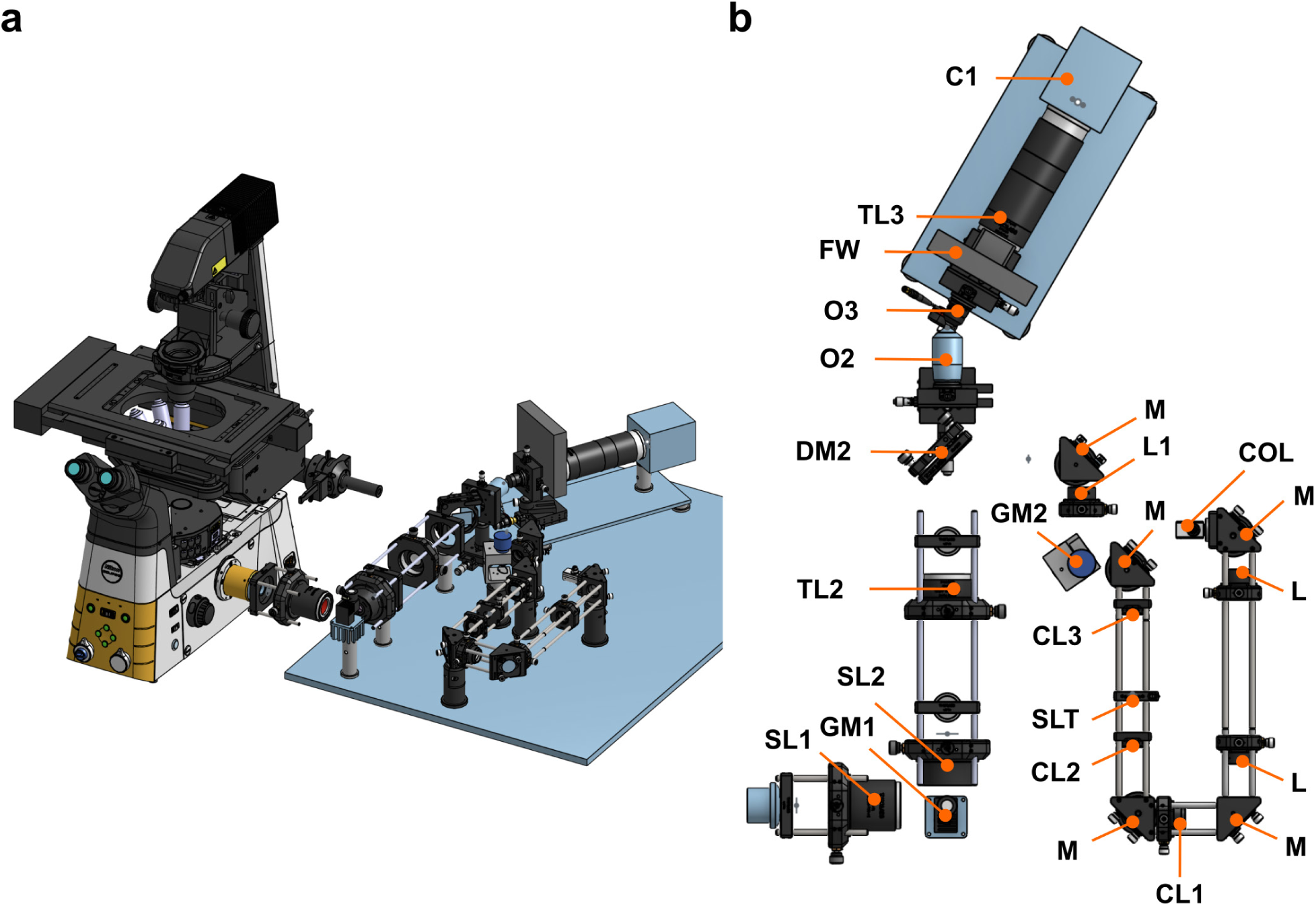
Mechanical layout of the remote refocus path for oblique plane microscopy. (a) Perspective view and (b) top-down view of a CAD model of the light-sheet remote refocus path. Symbols in panel (b) are as described in Figure 1-Supplement 1, with the addition of M: mirror, L: lens, COL: collimator.

**Figure 1-Supplement 3.**
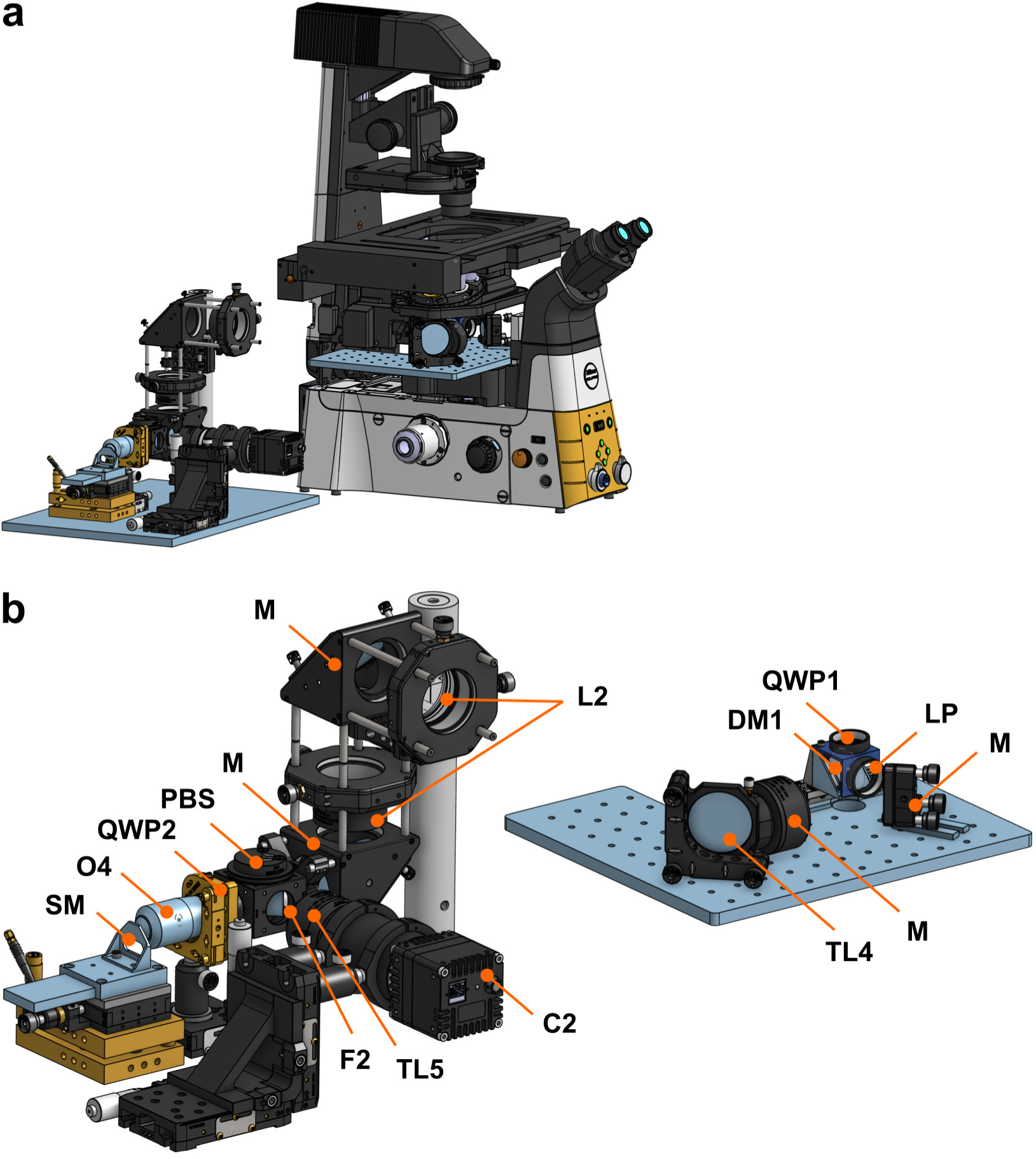
Mechanical layout of the remote-refocus path for label-free microscopy. Symbols in panel (b) are as described in Figure 1-Supplement 1 and Figure 1-Supplement 2.

**Figure 1-Supplement 4.**
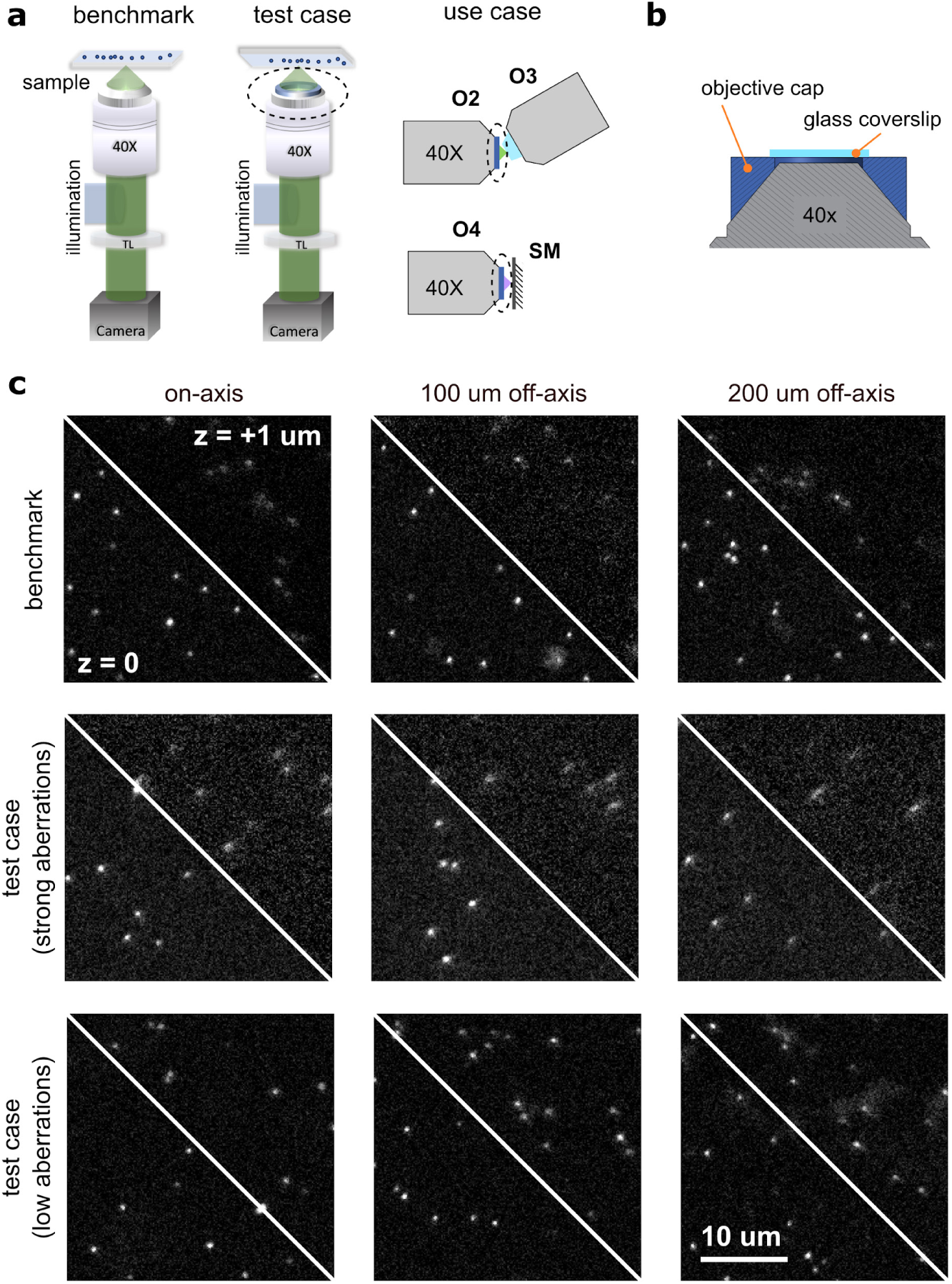
Attaching a glass coverslip to objectives O2 and O4. Objectives O2 and O4 are designed to image samples mounted on a #1.5 glass coverslip. As we are not using these objectives in the conventional manner, we needed to attach a coverslip to each objective to achieve aberration-free image formation of the remote volumes. We observed that tilt or strain in the glued coverslip can also introduce optical aberrations and therefore designed an iterative approach to adjust the coverslip angle and evaluate optical performance before the glue sets. We used a flexible adhesive to minimize strain (Dow DOWSIL 734) and analyzed the point-spread function of sub-resolution fluorescent beads to evaluate aberrations across the field of view. (a) Schematic of the optical setup used to characterize aberrations. Left: Experimental setup used to benchmark the performance of the objectives before gluing a coverslip; 100 nm fluorescent beads (Thermo Fisher Scientific, T7279) were dried on coverslip and imaged with epi-illumination. Middle: Experimental setup used to test for optical aberrations after a coverslip is glued. The sample is mounted upside-down with the beads facing the objective; a glass coverslip is attached to the objective using an objective cap we designed (highlighted with dashed line). Right: Intended use case of the objectives with glued coverslip in the remote refocus arms of the microscope. (b) Cross-section schematic of the objective cap. The objective cap positions the coverslip ∼100 µm away from the objective and allows us to attach the coverslip to the objective without damaging it. (c) Representative images showing bead point-spread functions for three cases: top, our benchmark case acquired following the left schematic in panel (a), middle, a test case showing poorly glue coverslip displaying coma aberrations across the field of view, acquired following the middle schematic in panel (a), and bottom, a test case showing correctly glued coverslip. Images are shown for in-focus (z = 0) and defocus (z = +1 um) planes. Aberrations are worse away from the optical axis-we show images on-axis (left column), 100 µm off-axis (middle column), and 200 μm off-axis (right column). Our iterative alignment process started by adjusting the objective cap on the objective. We mounted a glass slide on the microscope stage and secured it tightly. We applied glue to the top side of the objective cap and attached a high-precision coverslip (Thorlabs, CG15NH1) using tweezers. The objective cap was then positioned on the objective and the assembly was slowly moved upwards until the cap comes in contact with the glass slide and is leveled. At this point we removed the glass slide and imaged a sample with fluorescent beads. We compare the bead images with the benchmark images acquired before gluing the coverslip to evaluate the optical performance. If we did not observe strong aberrations, we placed the glass slide on the sample stage again and brought the objective cap in contact with the glass slide. We then carefully applied glue between the cap and the objective body with a fine pipette tip. We again evaluated for aberrations using the bead sample. If we were not satisfied we have the option to remove the glue from the objective cap and redo the process with a fresh coverglass piece. Case 2 in panel (c) shows data from one of our successful gluing procedures.

**Figure 1-Supplement 5.**
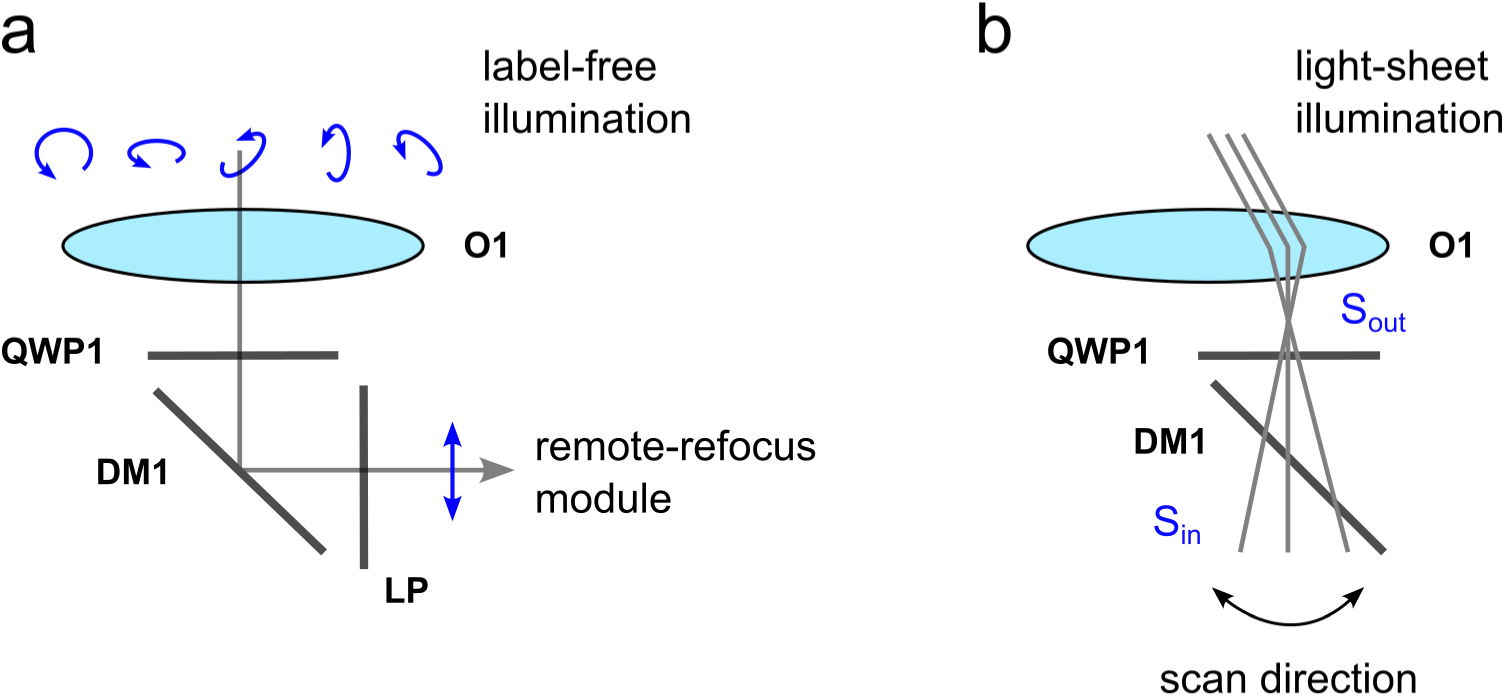
Design of the label-free analyzer cube. (a) The label-free analyzer cube was designed to convert physical properties of the sample encoded in the polarization of the transmitted light into intensity which will be detected by the camera. The sample is illuminated with right-hand circular or elliptical polarization and the analyzer cube is designed to work as a left-hand circular analyzer. In the absence of dichroic mirror DM1, quarter-wave plate QWP1 and linear polarizer LP act as a left-hand circular analyzer when the slow axis of QWP1 is oriented at 45 degrees from the transmission axis of LP. The dichroic mirror DM1 introduces an unknown retardance, which makes the QWP1-DM1-LP assembly deviate from a left-hand circular analyzer unless LP’s transmission axis is perpendicular to DM1’s slow axis. To make the assembly insensitive to DM1, we rotated QWP1 and LP together, maintaining their relative 45 degree orientation, until the complete optic behaves like a left-hand circular analyzer by extinguishing right-hand circular light. The key idea is that the behavior of a linear polarizer is unchanged by the addition of a retarder with a perpendicular slow axis. This is demonstrated by the Mueller matrices below, where *δ* is an arbitrary retardance. (b) The polarization of the light-sheet illumination beam is also changed as it travels through the cube. We have not intentionally controlled the polarization of the light-sheet illumination, and we measured its Stokes state before and after the cube to be S_in_ = [1.00, 0.48,-0.82, ±0.31] and S_out_ = [1.00,-0.75, 0.03, ±0.66], respectively, assuming full polarization of the beam. The schematic further illustrates the light-sheet scan direction relative to the tilt axis of DM1. 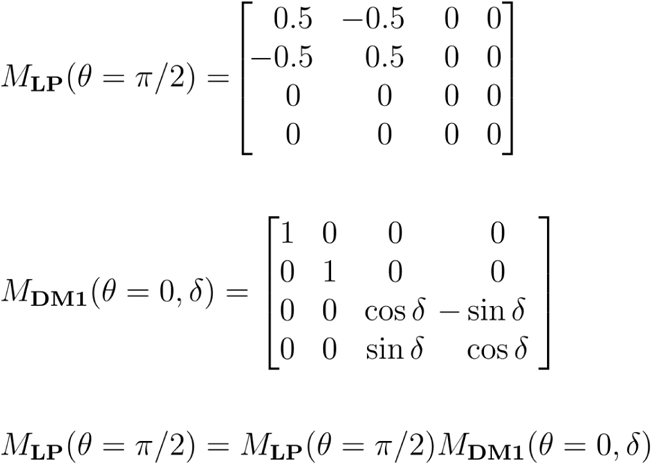

**Figure 1-Supplement 6.**
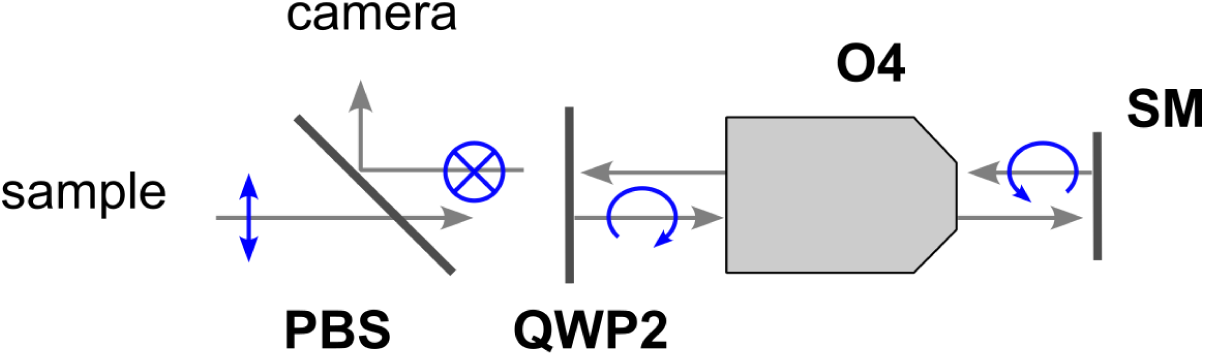
Design of the label-free polarized light filter cube. The filter cube is designed to maximize the throughput of light by modulating its polarization (blue symbols). Light from the sample is linearly polarized by LP in the analyzer cube and is transmitted by the polarizing beamsplitter PBS. We placed QWP2 behind O4 to create left-handed circular polarization. The polarization of light switches to right-hand circular upon reflection at the scan mirror. As the light passes through QWP2 again its polarization is converted to linear and orthogonal to the input beam. The beam is now reflected by PBS and directed towards the camera.

**Figure 1-Supplement 7.**
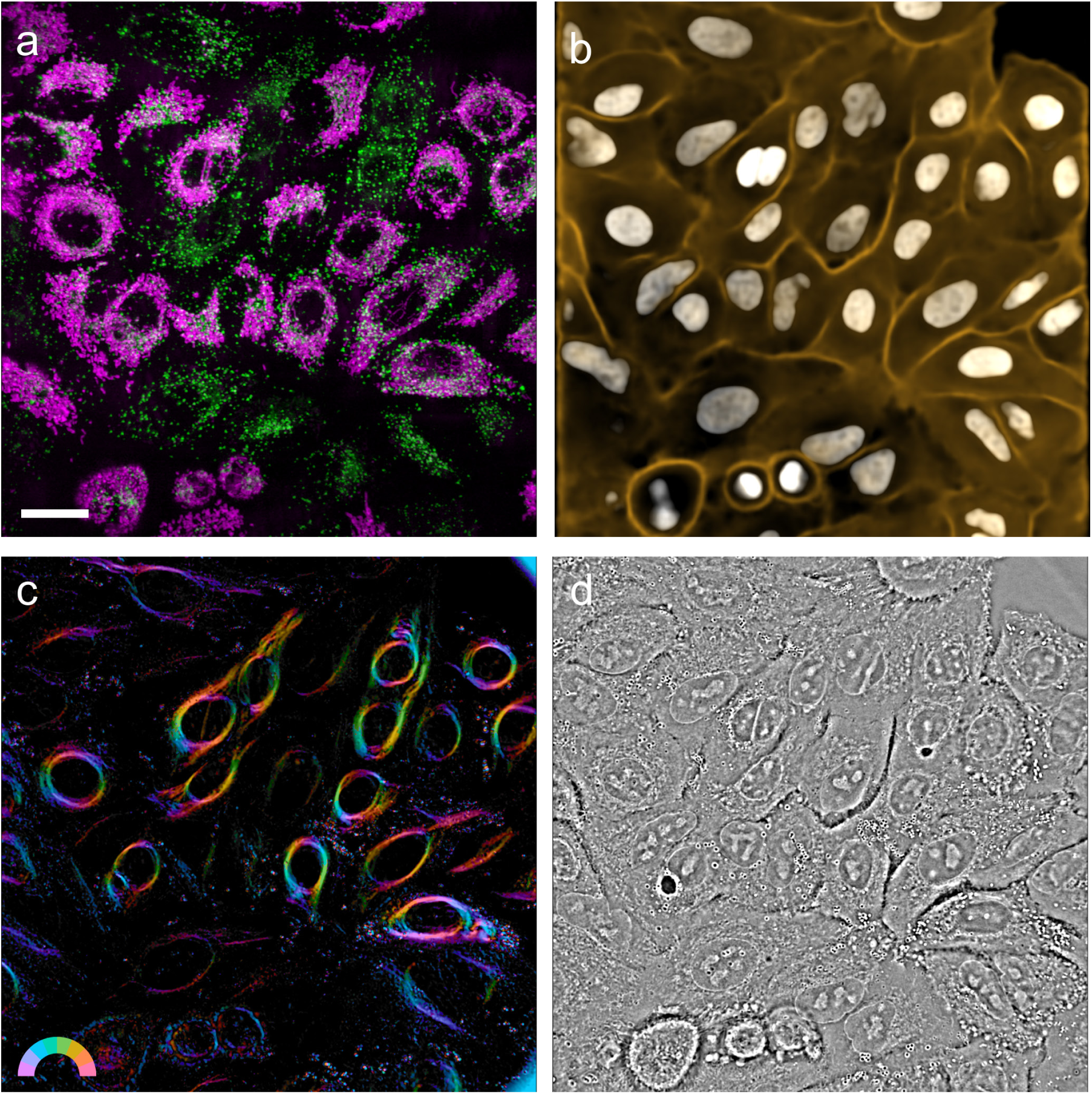
Full field of view display of the available imaging modalities. (a) Overlay of maximum intensity projection of labeled lysosomes (green) and mitochondria (magenta). (b) Overlay of mean intensity projection of predicted nuclei (gray) and membrane (orange). (c) Retardance and orientation overlay of a central axial slice. (b) Phase image of a central axial slice. Scale bar: 20 µm.

**Figure 1-Supplemental Note 1.**
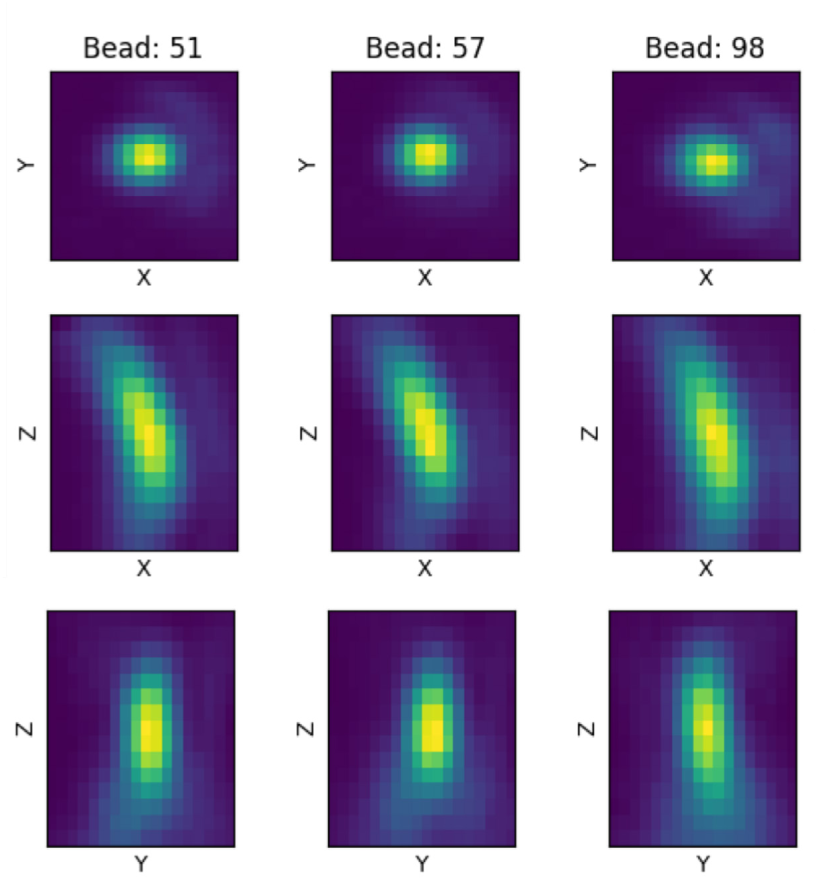
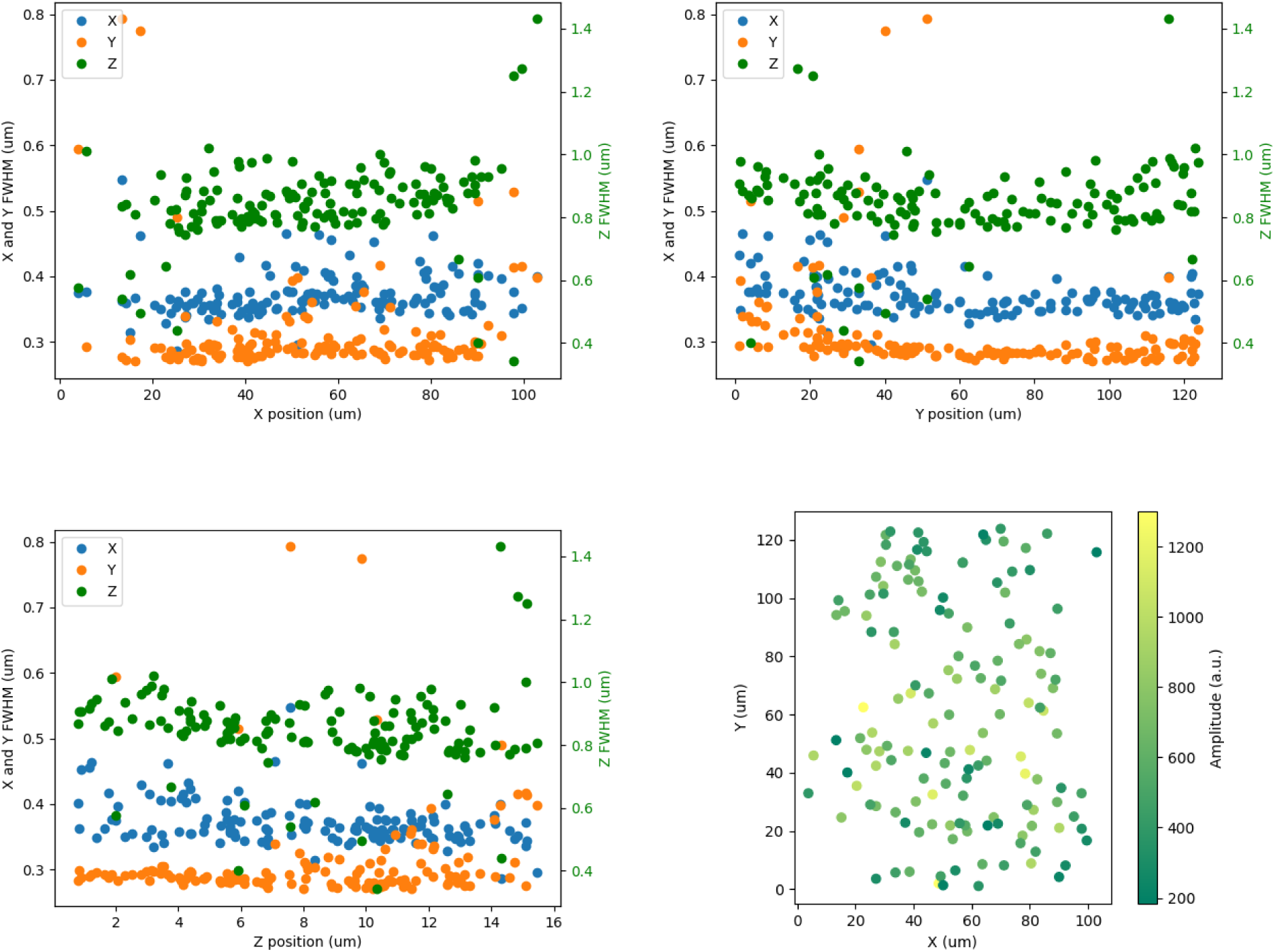
PSF benchmarking routine. We developed a fast online PSF benchmarking routine (available in https://github.com/czbiohub-sf/shrimPy) which was critical for optimizing the alignment of the microscope prior to experiments. The benchmarking routine consists of the following steps:

1. Collect 3D volume of beads
2. Detect beads in 3D
3. Optional: Deskew raw data
4. Measure the bead PSF full width at half maximum (FWHM) along all three axes
5. Generate an HTML report Here is a sample of the results from the generated report: **FWHM**

- 3D Gaussian fit
  X: 0.372 ± 0.128 um
  Y: 0.311 ± 0.075 um
  Z: 0.845 ± 0.034 um

- 1D profile
  X: 0.296 ± 0.036 um
  Y: 0.262 ± 0.012 um
  Z: 0.585 ± 0.010 um

- 3D principal components
  ○ 0.284 um, 0.332 um, 0.875 um We found it critical to use beads dispersed in media with a refractive index of 1.40, matching the design specifications of the microscope. We developed a protocol for embedding beads in 0.5% low melting point agarose (Sigma-Aldrich A2576) dissolved in 50% (w/w) glycerol in water (53). We found this method to be superior to using beads embedded in Fluoromount-G or agarose dissolved in water, which showed increased PSF FWHM as a function of depth through the sample. It was also very important that this PSF benchmarking routine is fast enough to be performed repeatedly during microscope alignment. We developed GPU-accelerated methods for detecting beads and deskewing raw data. A typical iteration of acquiring and analyzing data takes approximately 30 seconds. To characterize the bead PSF we used methods available in napari-psf-analysis. We report FWHM in three dimensions based on: (1) 3D Gaussian fit to the bead volume, (2) 1D Gaussian fit to slices through the center of the bead volume, and (3) principal axes given by the eigenvectors of the fitted 3D Gaussian covariance matrix. In cases where the measured PSF deviates from a Gaussian profile we report the 1D FWHM measurements. We find that accurate FWHM measurements require fine sampling of the PSF profile, so we used an Oryx ORX-10GS-123S6M-C (Teledyne FLIR, 3.45 µm pixel size) camera instead of the Prime BSI Express camera (6.5 µm pixel size) during alignment. We use this benchmarking routine prior to experiments to set the O1 correction collar for the given temperature and sample coverslip thickness. We also use this benchmarking routine for troubleshooting the microscope alignment, for example in a straight (i.e. 0 degree) configuration between O2 and O3, with O2 and O3 removed from the optical path, or using epi-rather than light-sheet illumination.

**Figure 1-Supplemental Movie 1.**
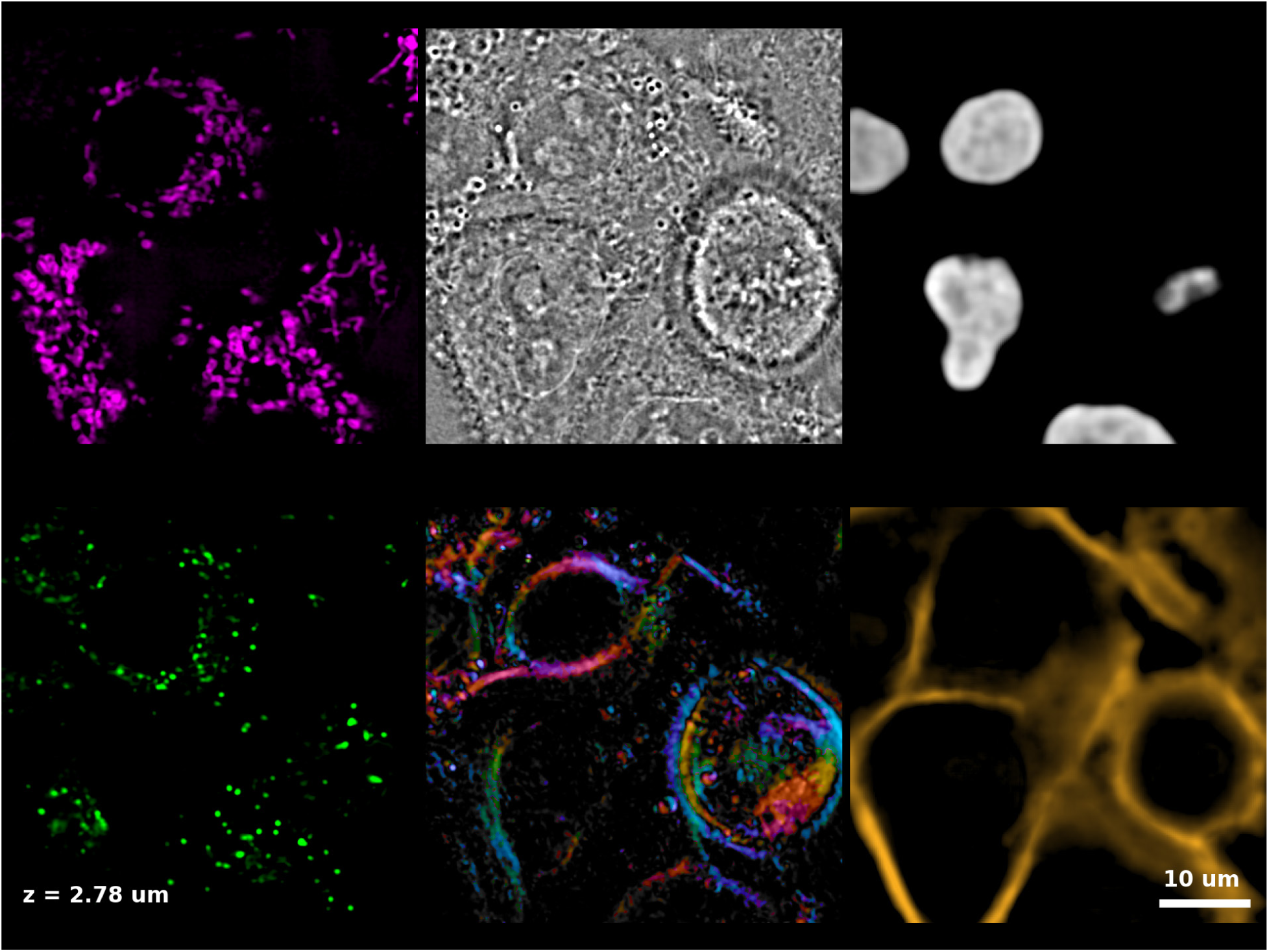
Axial fly-through of A549 cells in all imaging modalities. Top row: labeled mitochondria, phase, and nucleus prediction. Bottom row: stained lysosomes, birefringence, and membrane prediction. The same region of interest as in Figure 1b is shown.

**Figure 1-Supplemental Movie 2.**
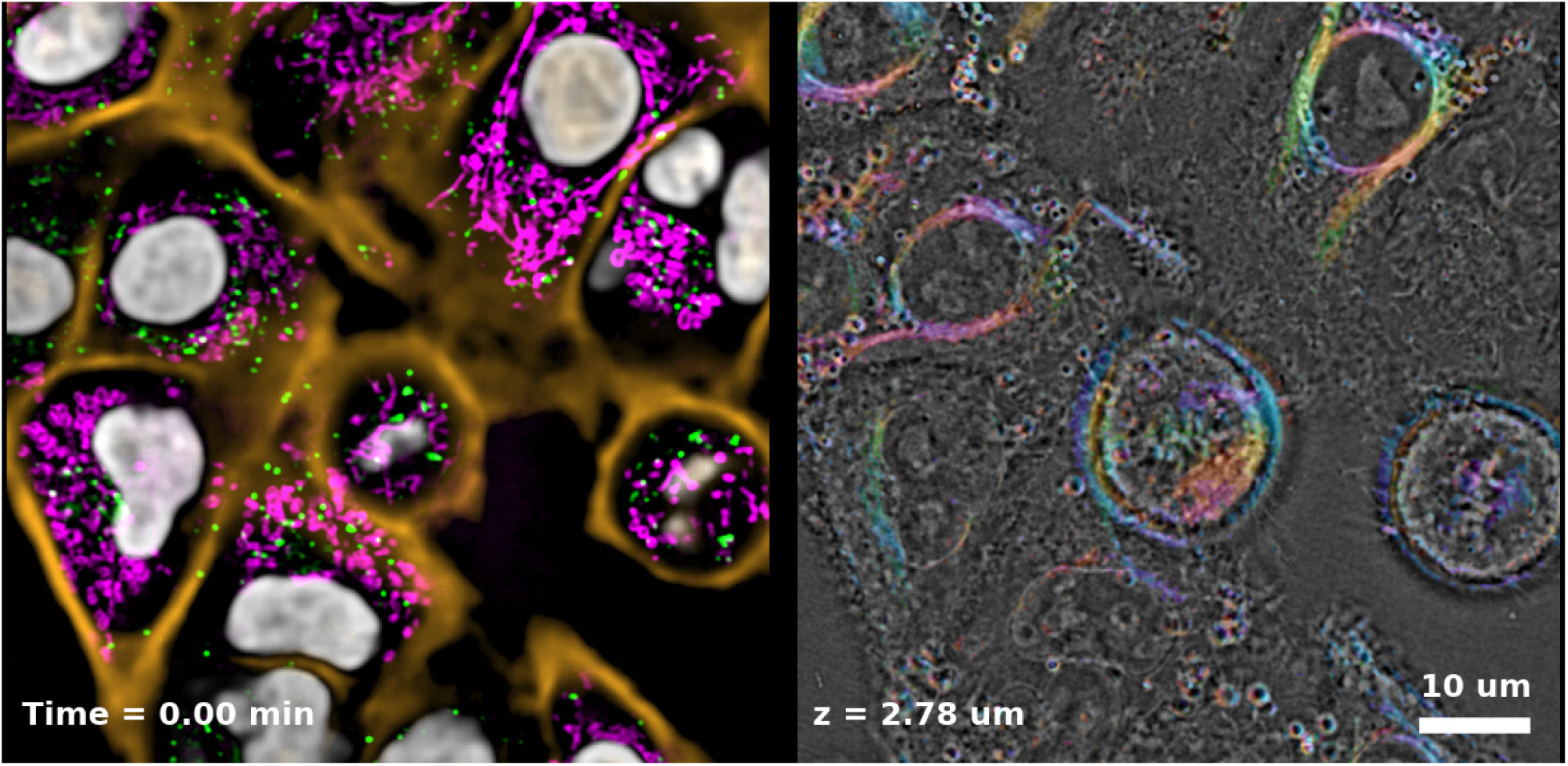
Timelapse of A549 cells in all imaging modalities. Left: overlay of mitochondria (magenta), lysosomes (green), nuclei (gray), and membrane (orange). Right: overlay of birefringence (color) and phase (grayscale). Data are acquired at 2 volumes per minute.

**Figure 1-Supplemental Movie 3.**
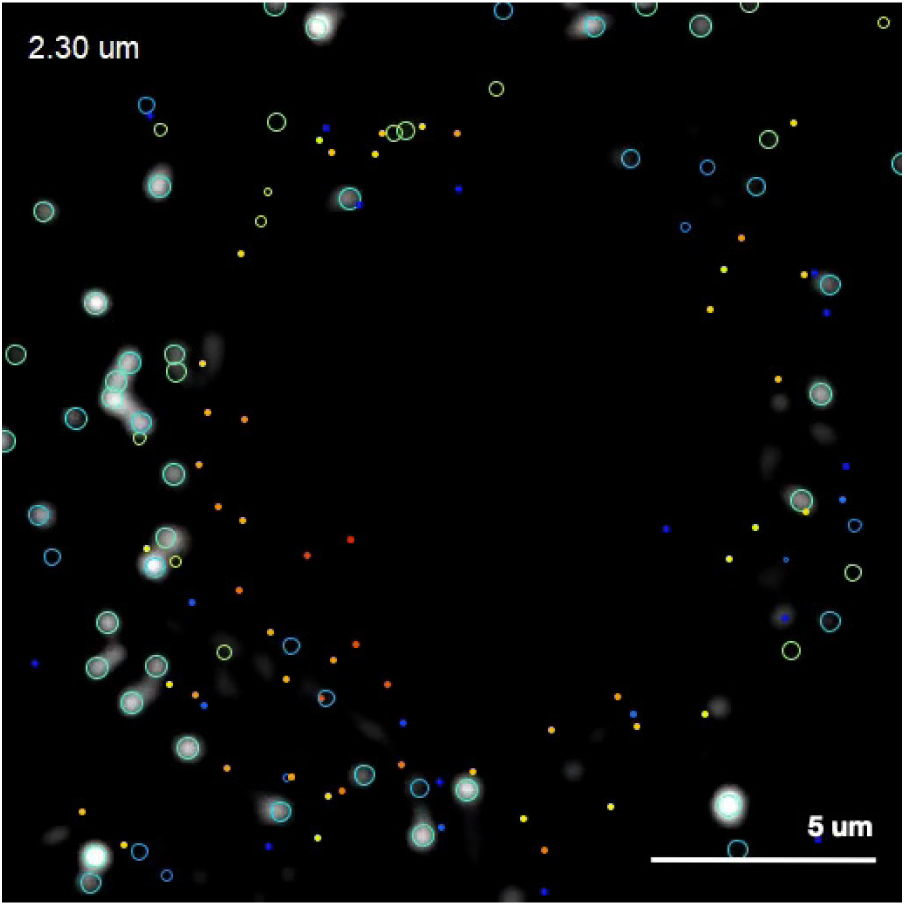
Tracking of lysosomes in A549 cells. Spots and tracks are color-coded by depth as in Figure 1d. Circles show in-focus spots, dots show out-of-focus spots. The timelapse shows tracks over a smaller 2.3 µm axial section of the data.

**Figure 2-Supplement 1.**
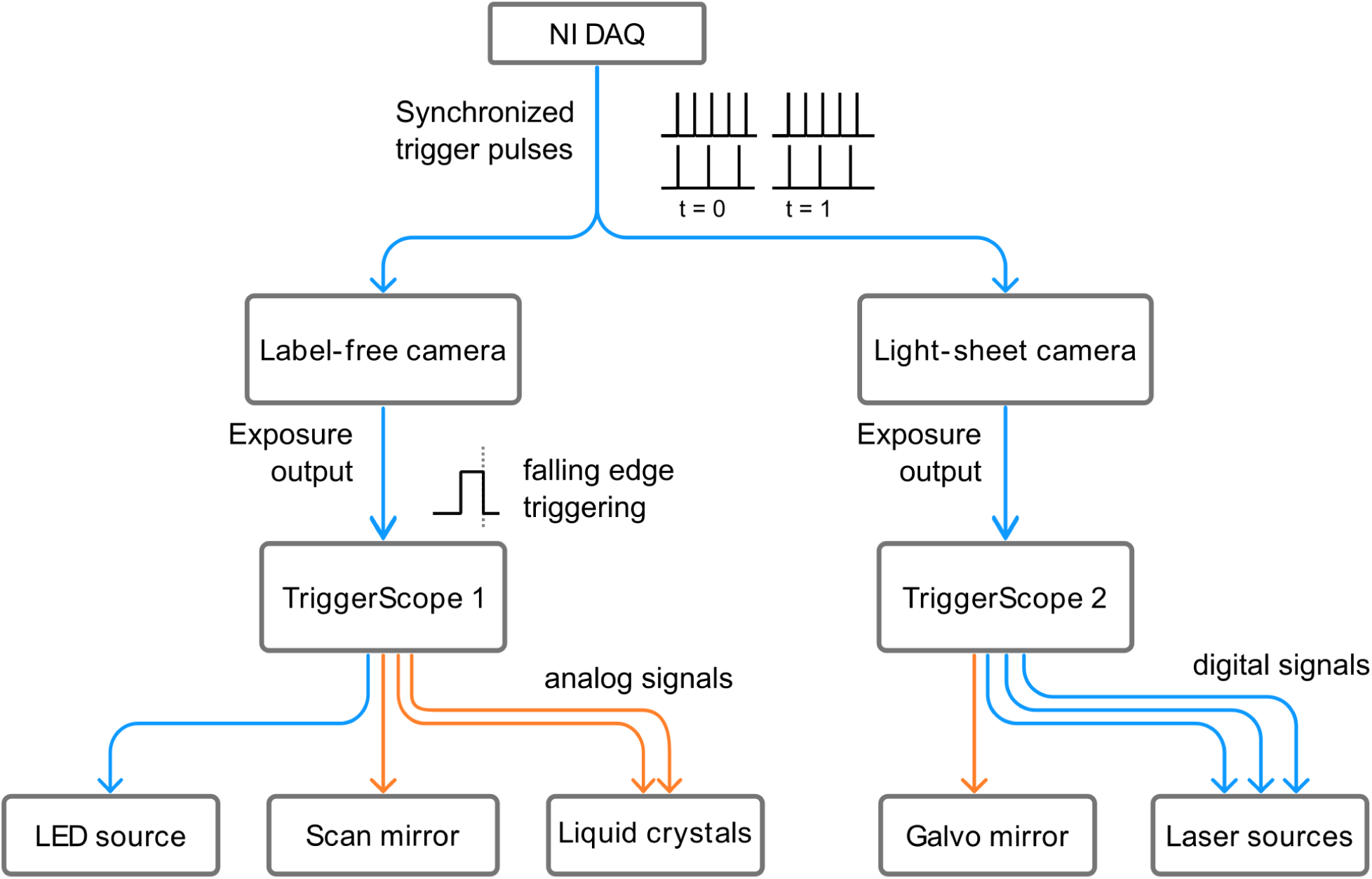
Hardware synchronization diagram. Acquisition on the Mantis microscope is synchronized via analog and digital signals. For each timepoint the shrimPy acquisition engine generates a set of synchronized trigger pulses on a National Instruments DAQ card which are routed to the two cameras; the label-free and light-sheet acquisitions typically have a different number of channels and z-slices. The exposure output of each camera is connected to the input of a dedicated TriggerScope which sends analog (orange) and digital (blue) signals to the associated hardware. Microscope hardware is changed on the falling edge of the exposure output pulse, in preparation for the following frame.

**Figure 2-Supplement 2.**
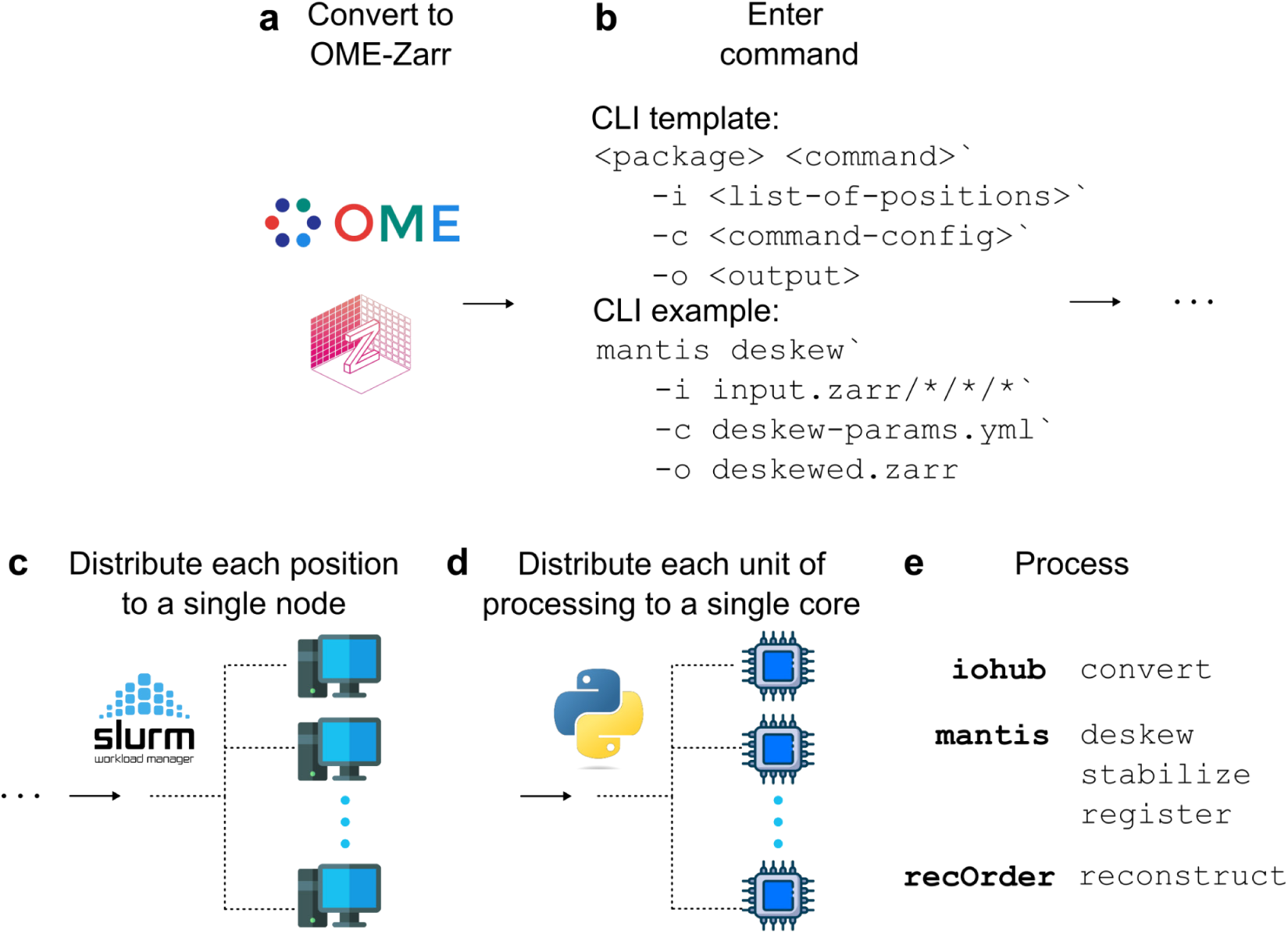
Parallel compute workflow. (a) We convert each dataset into OME-Zarr arrays in HCS (high-content screening) format with TCZYX dimensions, and (b) we apply processing operations by providing a list of input paths, where each path corresponds to a position, through a command-line interface (CLI). Our CLI uses the Slurm job scheduler to (c) distribute each position to a single node, and (d) Python multiprocessing further distributes each unit of data, either ZYX or CZYX arrays, for processing on single cores. (e) Our internal libraries use uniform CLI calls, enabling rapid prototyping and concurrent processing of multi-terabyte datasets by hundreds of cores.

**Figure 2-Supplement 3.**
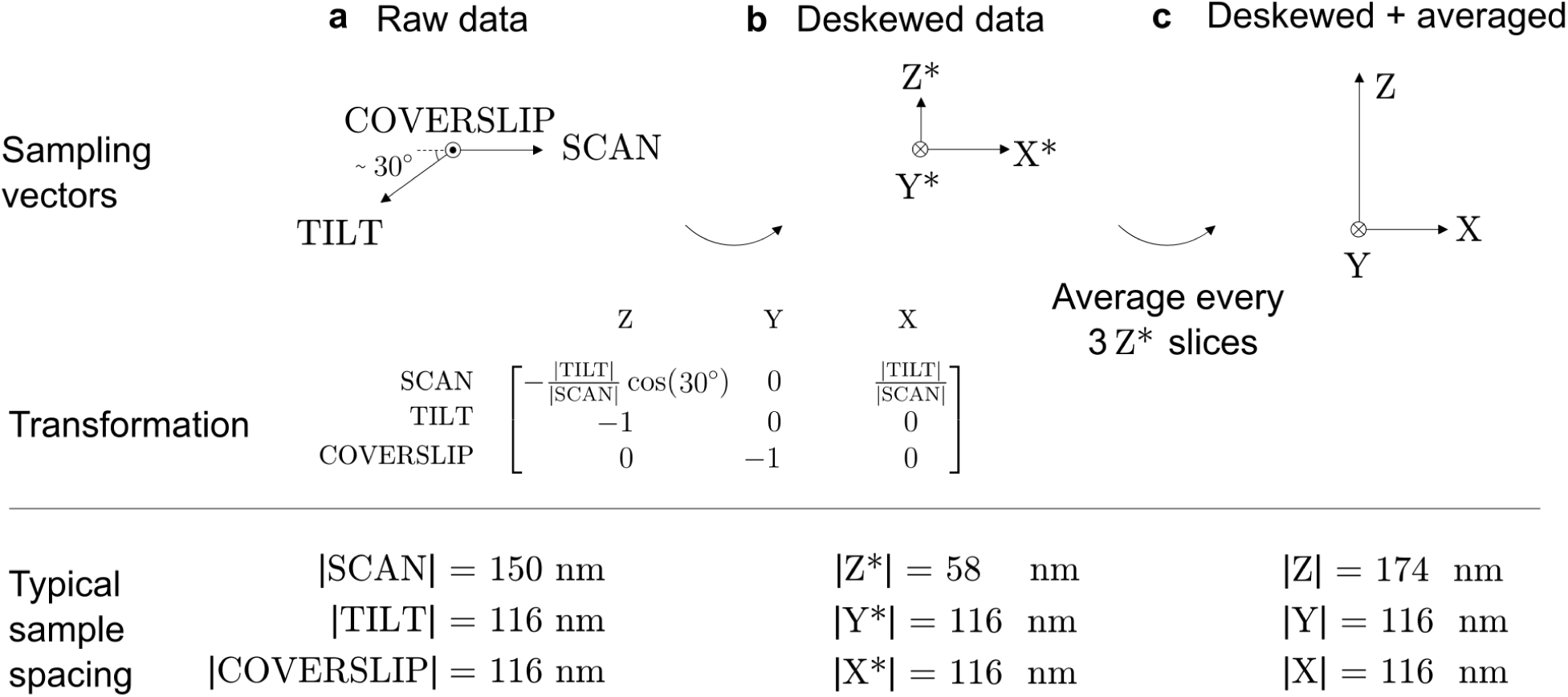
Deskewing workflow. We acquired raw data on a non-Euclidean sampling grid (a), which we deskewed by applying an affine transformation matrix (b), then averaged every three slices to improve SNR without sacrificing Nyquist sampling (c). The top row shows the sampling vectors, where the length of each vector indicates the distance between samples, the middle row shows the affine transformation matrix, and the bottom row shows our typical sample spacing. The oblique plane illumination and detection leads to oversampling of the dimension parallel to the optical axis of the objective (Z). For example, our microscope uses object-space pixels spaced by 6.5 *μ*m / (1.4×40) = 116 nm which correspond to Z-axis samples spaced by (116 nm) sin(30°) = 58 nm. This dimension is oversampled compared to our measured axial FWHM of ∼ 680 nm (Figure 2b, Figure 2-Supplement 4). Therefore, it is safe to average every three slices, boosting the signal-to-noise ratio and reducing data size without losing resolution.

**Figure 2-Supplement 4.**
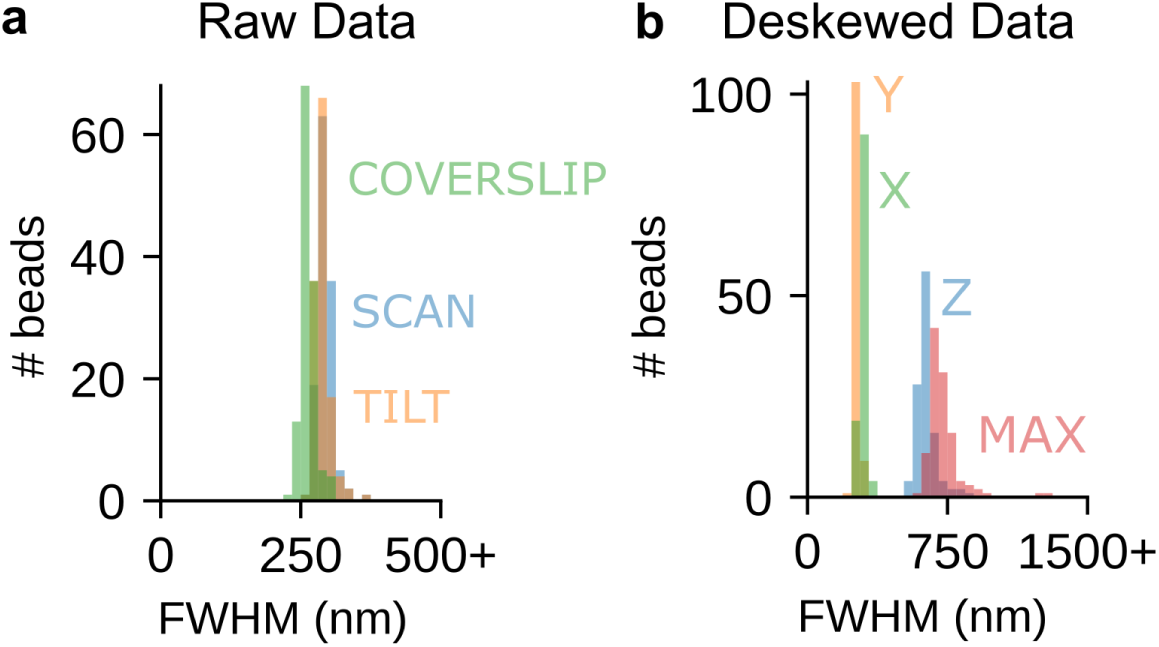
Light-sheet bead resolution measurements. We selected ROIs of 127 beads and measured FWHMs with respect to (a) raw sampling axes: SCAN = 294 ± 14 nm, TILT = 289 ± 14 nm, COVERSLIP = 263 ± 13 nm (mean ± std.), and (b) the deskewed sampling axes: Z = 683 ± 55 nm, Y = 262 ± 13 nm, X = 294 ± 15 nm. Additionally, we used the fluorescence intensities to calculate the moment-of-inertia tensor for each bead, and we used the tensor’s principal axis with the smallest moment of inertia to measure the longest axis of the PSF, labeled MAX FWHM = 726 ± 94 nm.

**Figure 2-Supplement 5.**
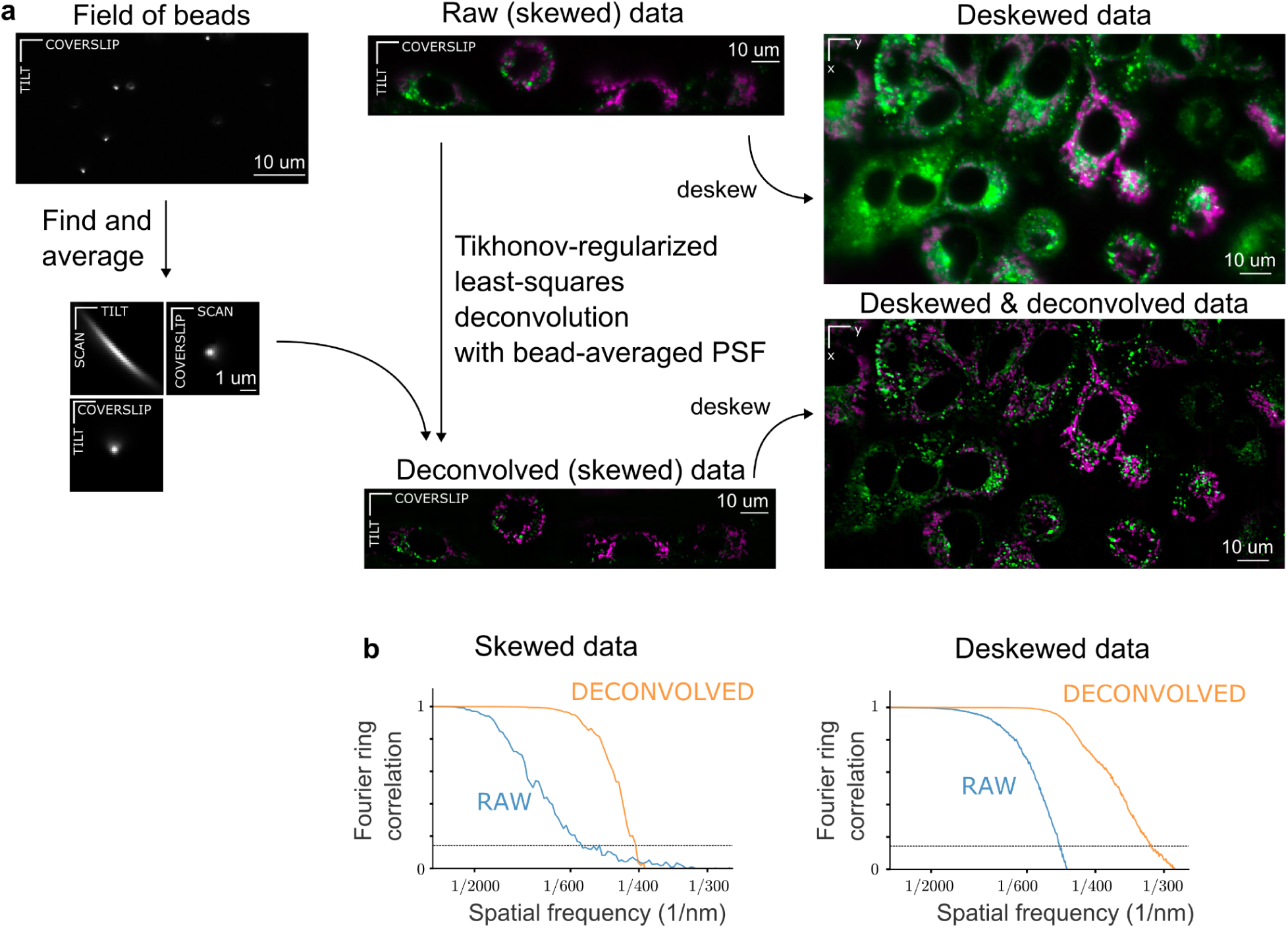
Fluorescence preprocessing workflow and Fourier ring correlation (FRC) resolution measurements. (a) Starting with a 3D volume of beads, we find and isolate bead patches, average them into a single PSF (assuming spatial shift invariance across the field of view), then use a Tikhonov-regularized least squares deconvolution to improve contrast and resolution (center column) before deskewing (right column). Here we show A549 cells with labeled mitochondria (TOMM20, magenta) and lysosomes (LysoTracker, green). In Figure 1 and associated movies we deskew and deconvolve (bottom right), while elsewhere we only deskew (top right) raw data. (b) Using the lysosome channel, we calculated Fourier ring correlation curves in skewed COVERSLIP-TILT coordinates (left) and deskewed X-Y coordinates (right). To calculate each curve we started with a single image, applied a ⅛ Tukey window, performed a pixel-wise binomial split, calculated the normalized radially averaged correlation of the two images’ spatial frequencies, then plotted the average of curves from 15 adjacent slices (Z slices for X-Y, SKEW slices for COVERSLIP-TILT). Before deconvolution (blue lines), we measured a smaller cutoff frequency (horizontal line), i.e. lower resolution, in COVERSLIP-TILT coordinates (∼1/550 nm^-1^) than in deskewed coordinates (∼1/500 nm^-1^) because principal axes of the PSF are more closely aligned with the deskewed X-Y axes. After deconvolution (yellow lines), the cutoff frequencies are increased to ∼1/400 nm^-1^ for skewed data and ∼1/300 nm^-1^ for deskewed data.

**Figure 2-Supplement 6.**
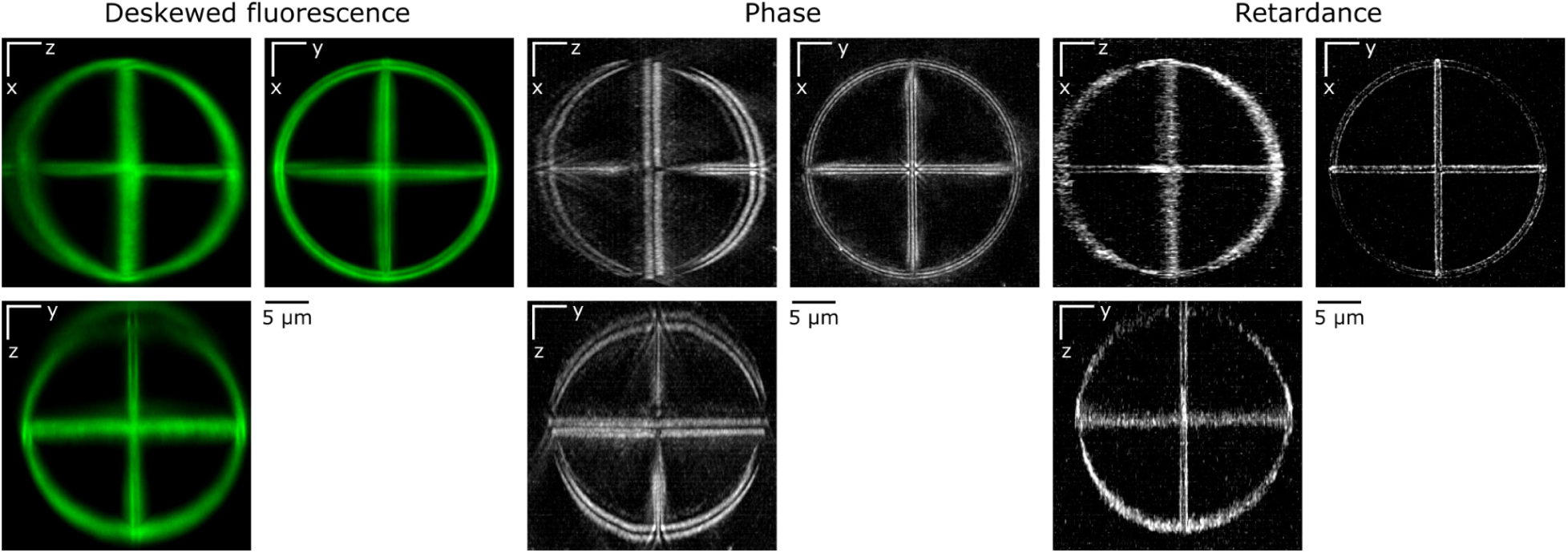
Maximum intensity projections of Argolight sphere in deskewed fluorescence (left), phase (center), and retardance (right) channels.

**Figure 2-Supplement 7.**
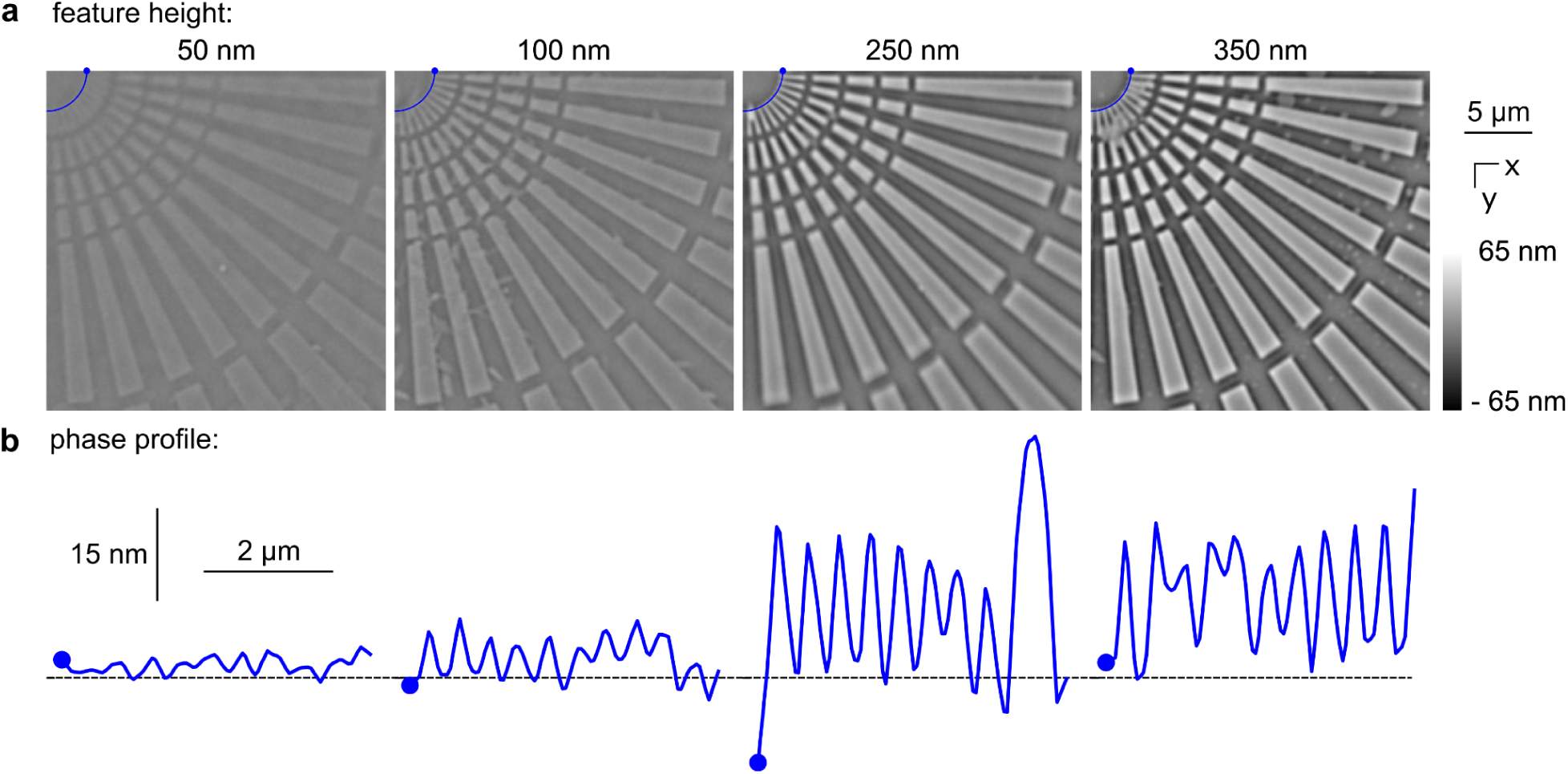
Phase resolution target. (a) We reconstructed the in-focus phase from a remote-refocus stack of a resolution target with glass feature heights of 50, 100, 250, and 350 nm (columns, left to right), and (b) we plotted line profiles through the spoke pattern’s minimally separated bars spaced by 400 nm at their narrowest points (near the top left corner of each image). We are able to resolve these bars for feature heights as small as 100 nm, and we begin to lose these signals in the noise for 50 nm feature heights.

**Figure 2-Supplement 8.**
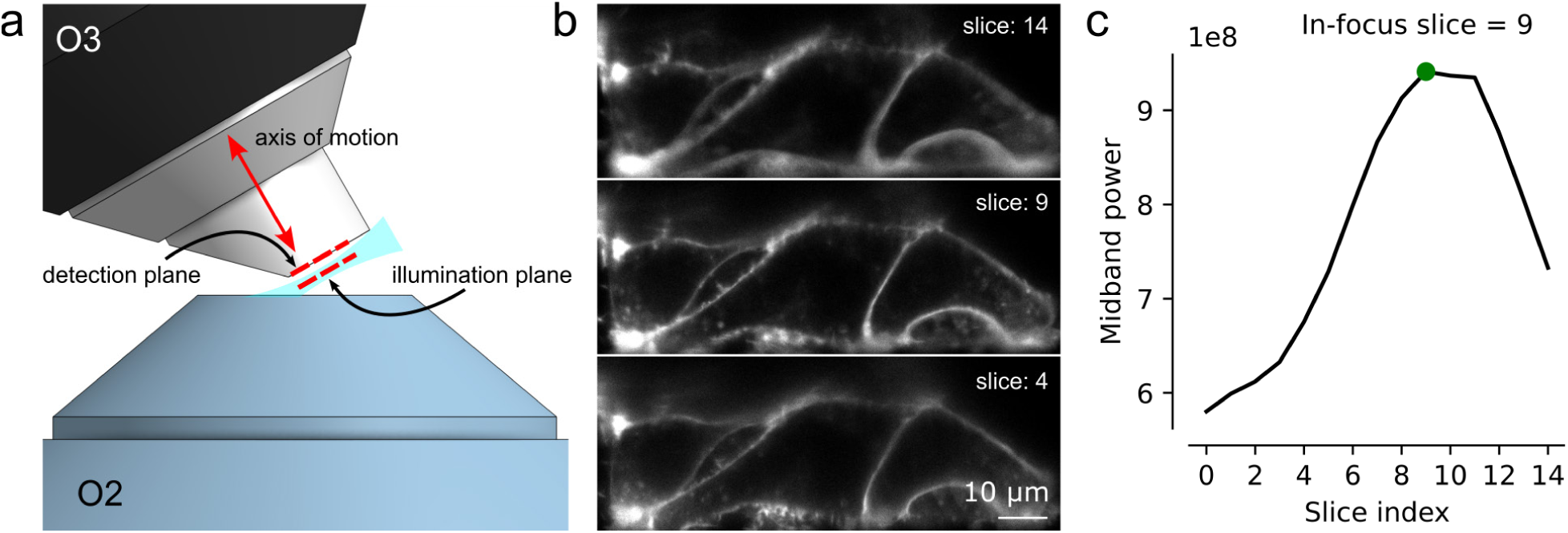
Light-sheet arm hardware autofocus. (a) The O3 objective is translated along its optical axis to align the plane illuminated by the light sheet with the detection plane of the O3 objective. Images are acquired at consecutive planes (slices) and analyzed to determine the optimal objective position. (b) Features in the sample are sharply in focus when the illumination and detection planes coalign. (c) The optimal O3 position is chosen by maximizing the image’s transverse midband power—the total power in the region *v*_c_/8<|*v|<v_c_*/4, where *v* is the 2D transverse spatial frequency and *v*_c_ = 2NA/ λ is the transverse cutoff frequency.

**Figure 2-Supplement 9.**
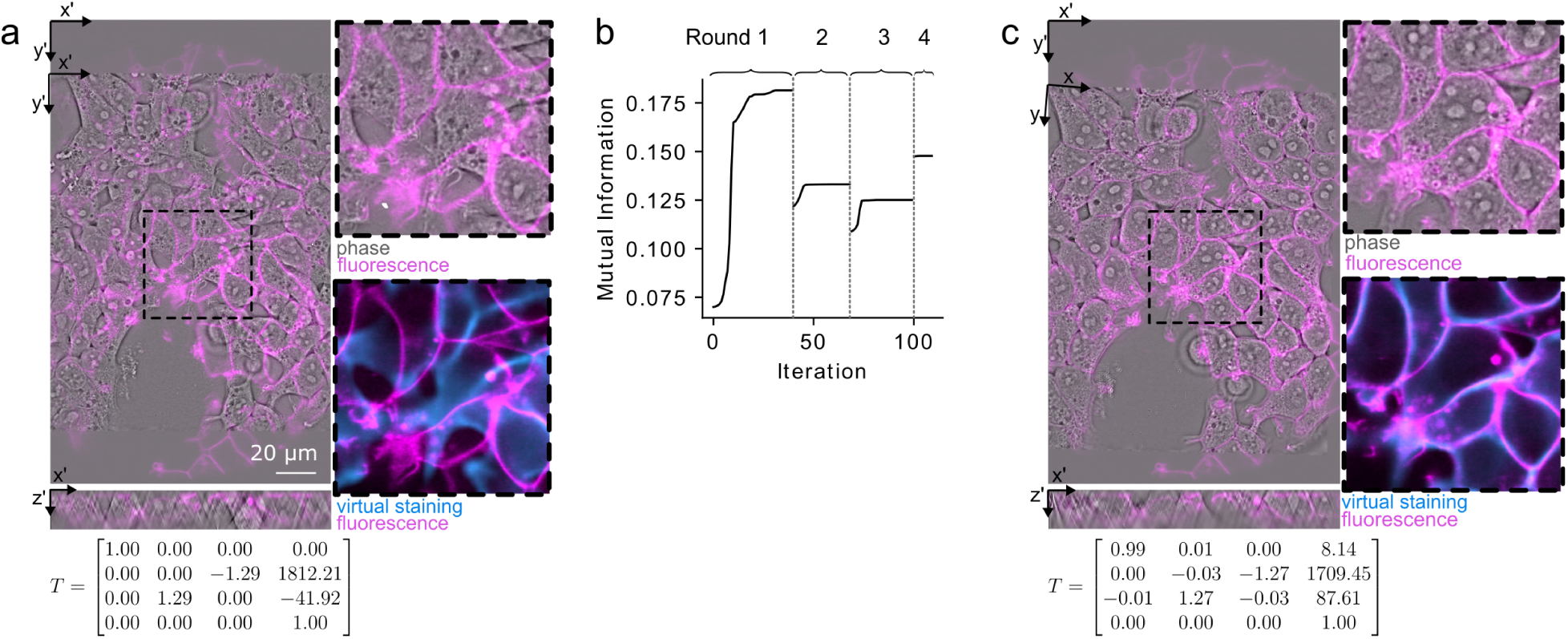
Registration workflow with virtual staining. (a) We used virtually stained organelles to register phase and fluorescence volumes. We do coarse registration interactively using napari to visualize the volumes (rotation of the phase volume by 90 degrees, rescaling the voxels to the same size, and rigid transformation in the XY plane based on the key points selected by a user). (b) The transformation is then refined by maximizing the mutual information between virtual staining and fluorescence volumes using four rounds of optimization. In the first round, both volumes are downsampled by 6x and the 3D similarity transform that maximizes mutual information is found. Subsequent rounds start with the result of the previous round with decreasing downsampling—the second, third, and fourth rounds use 4x, 2x, and no downsampling, respectively. (c) The optimized transformation matrix is applied to the phase volume. Transformation matrices shown in (a) and (c) are expressed in homogeneous ZYX-ordered voxel coordinates.

**Figure 2-Supplemental Movie 1.**
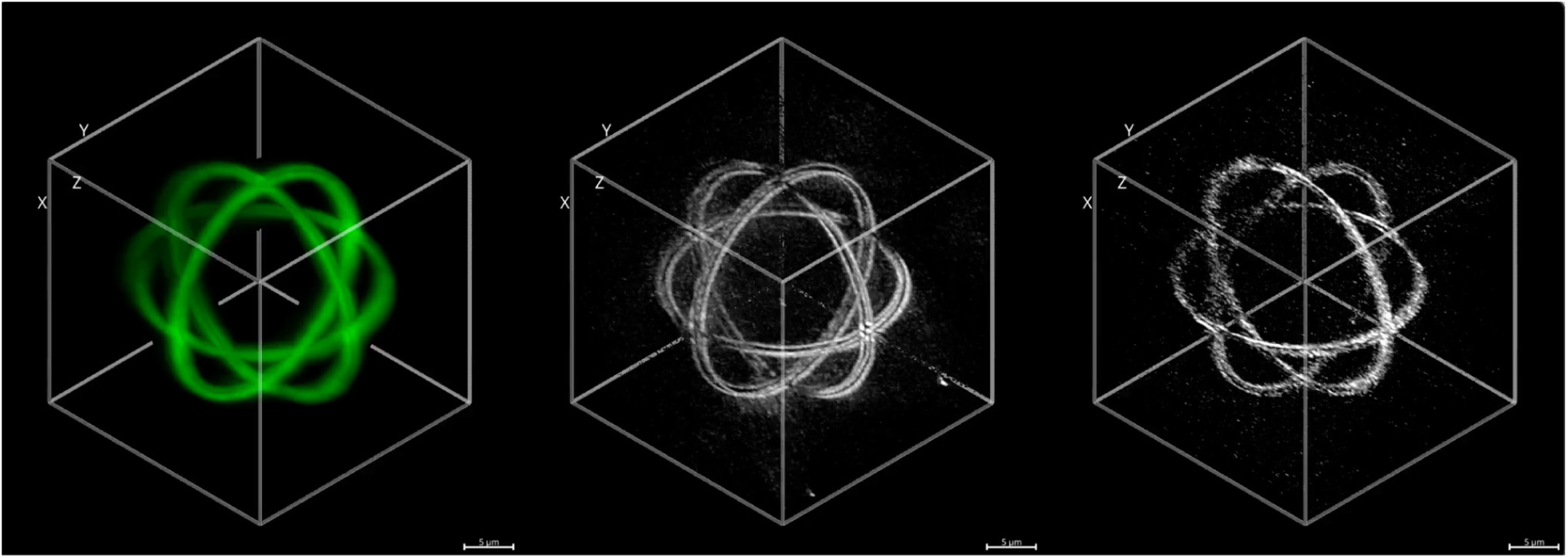
Fly around of Argolight sphere in deskewed fluorescence (left), phase (center), and retardance (right) channels.

**Figure 2-Supplemental Movie 2.**
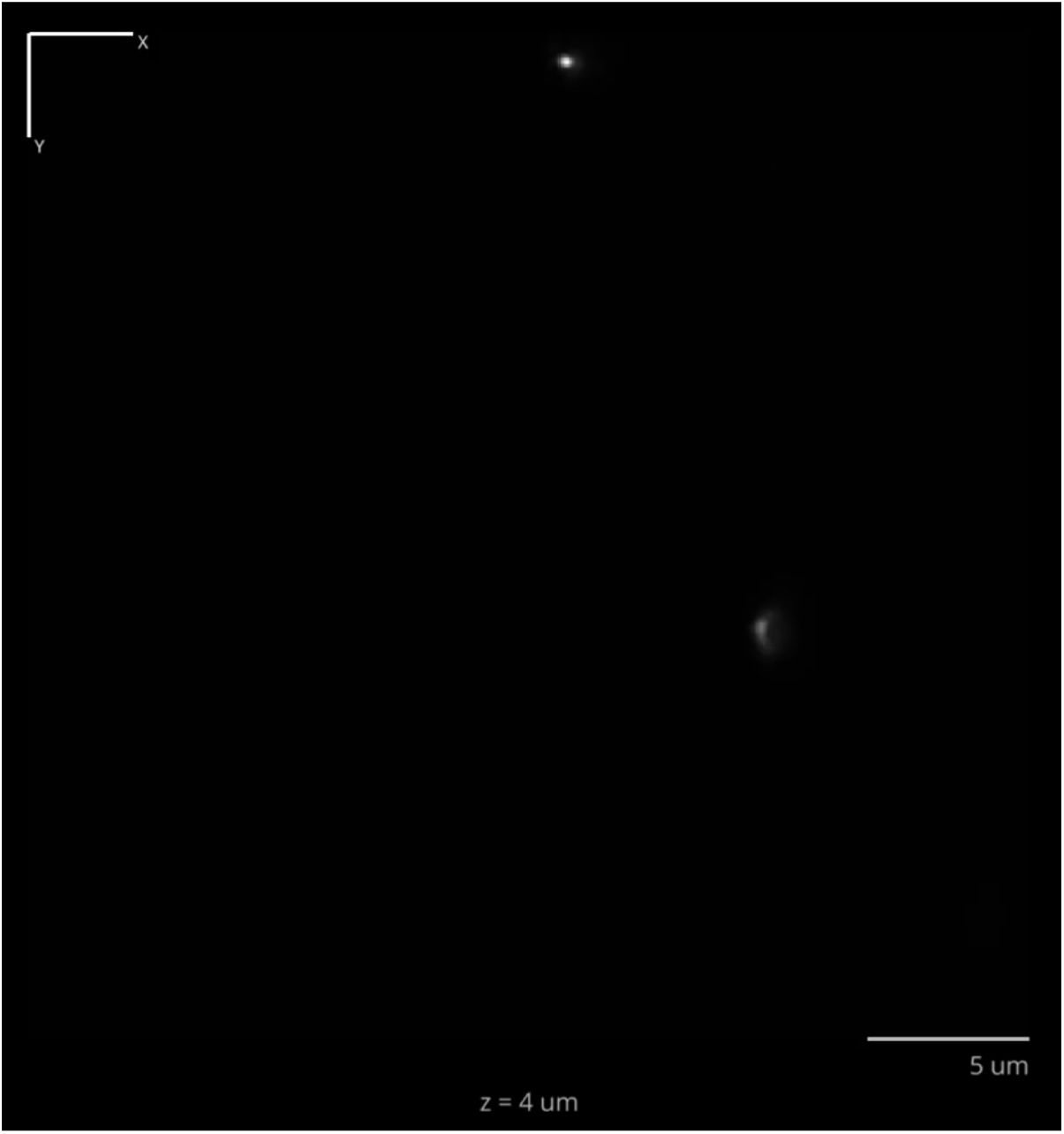
Axial fly-through of fluorescent bead PSFs in deskewed coordinates.

**Figure 3-Supplemental Movie 1.**
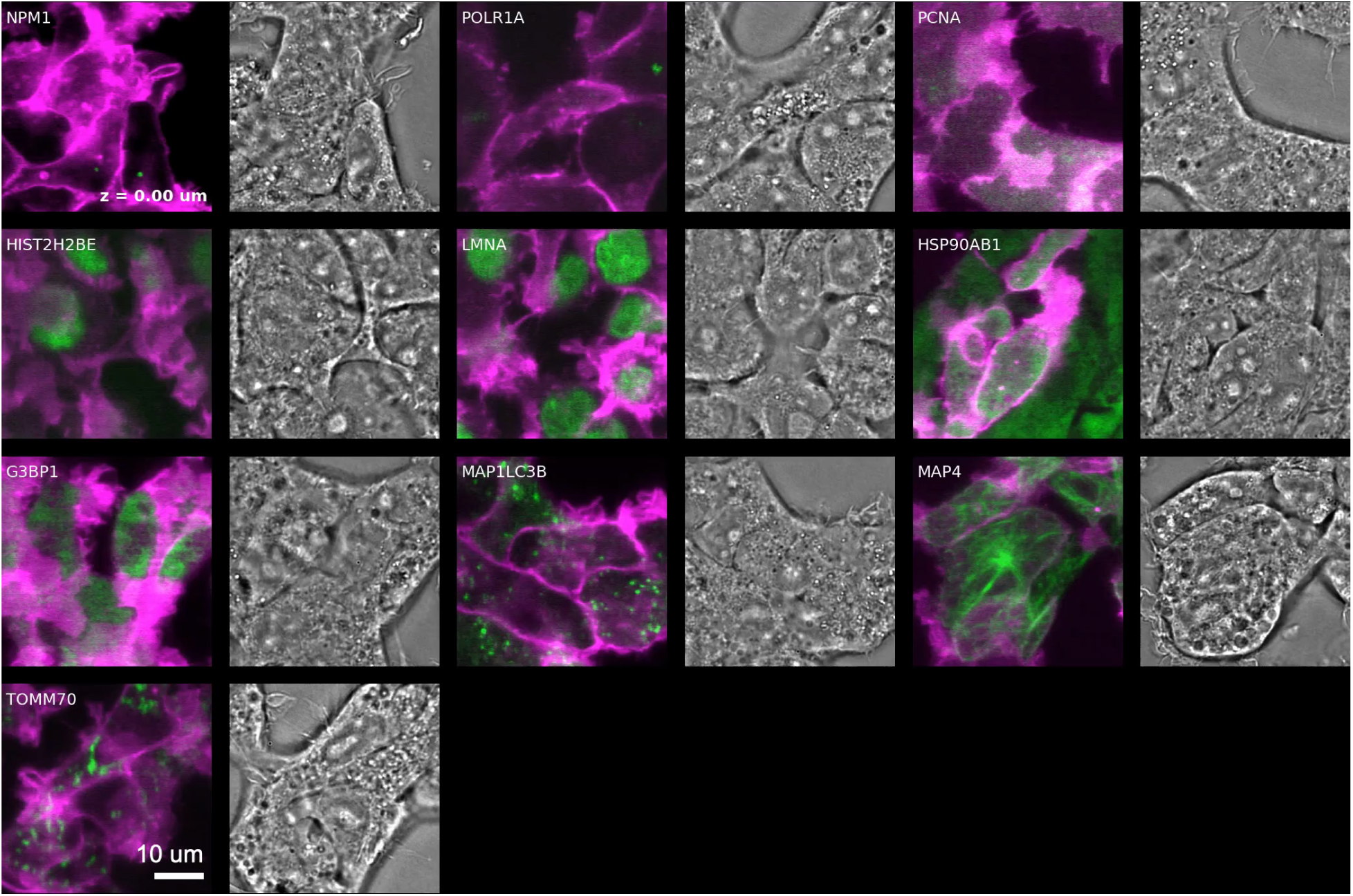
Axial fly-through of OpenCell targets 1-10. For each target we show a pair of images-overlay of molecular markers in green and magenta on the left, and phase image on the right.

**Figure 3-Supplemental Movie 2.**
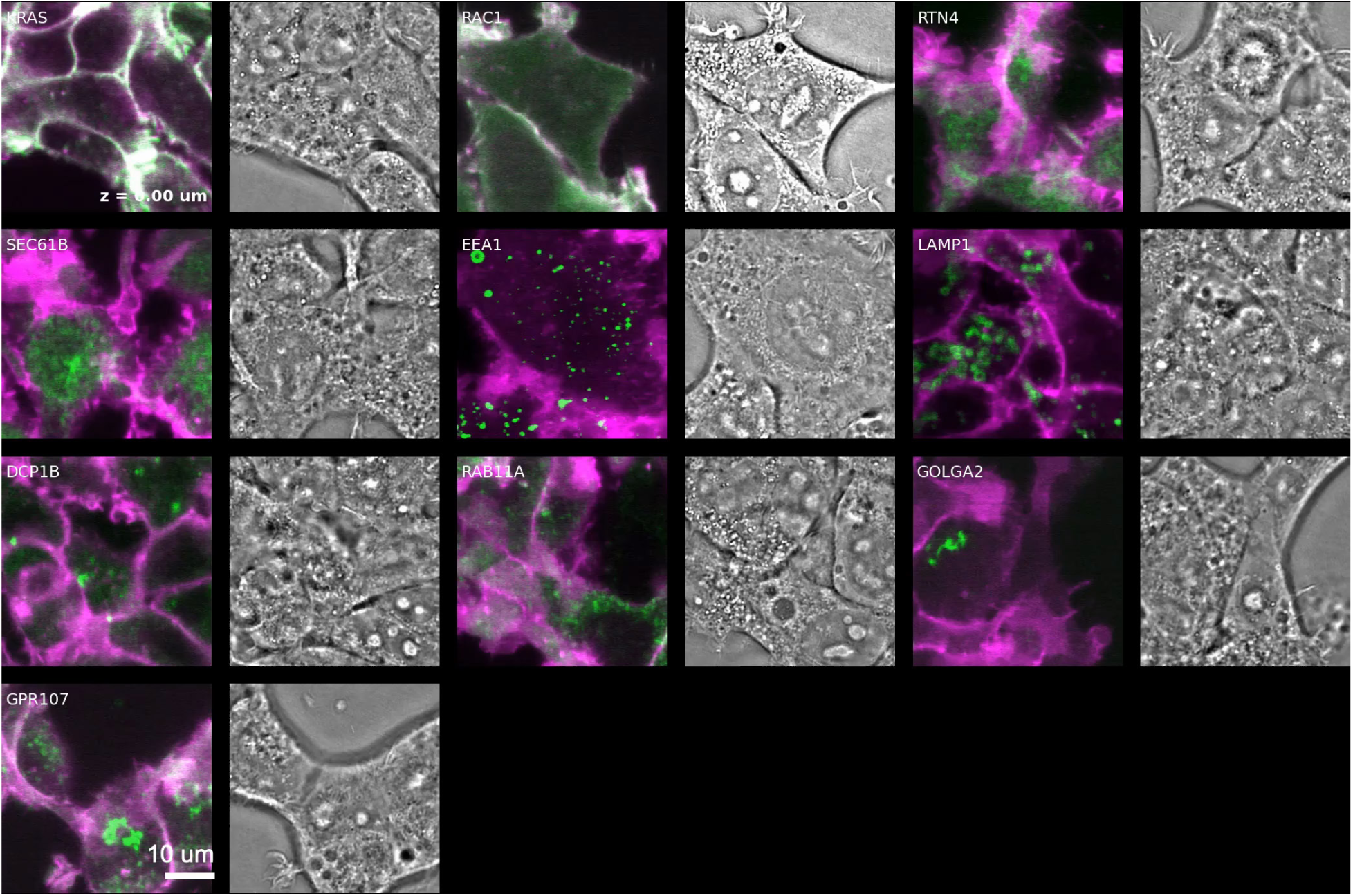
Axial fly-through of OpenCell targets 11-20 displayed as in Figure 3-Supplemental Movie 1.

**Figure 4-Supplement 1.**
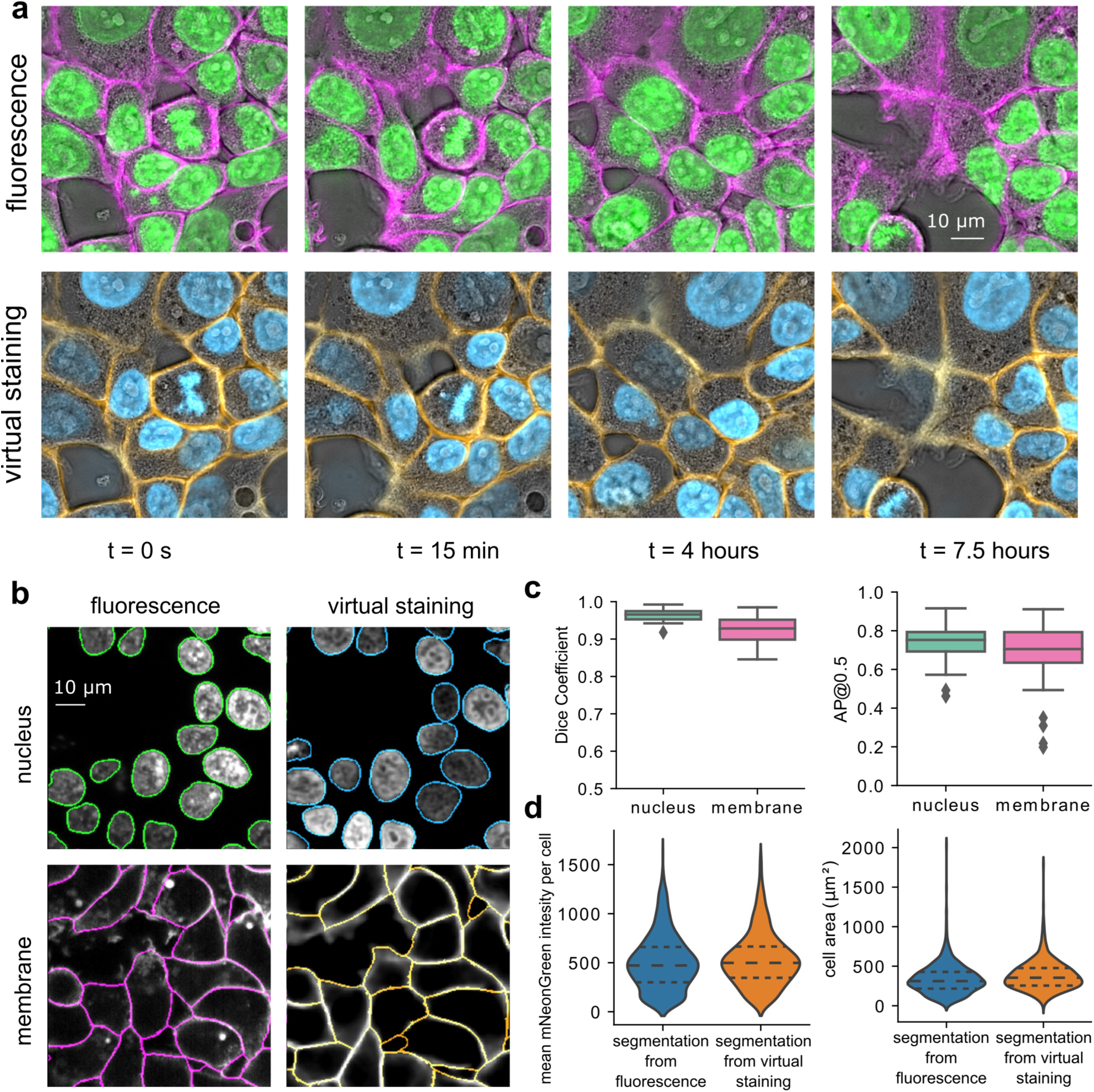
Evaluation of the virtual staining and segmentations. (a) Virtual staining of the nucleus and plasma membrane for the HIST2H2BE cell line expressing HIST2H2BE-mNeonGreen (green) and CAAX-mScarlet (magenta). Top row shows fluorescent channels, bottom row shows virtually stained channels. (b) Instance segmentation masks of the nucleus (top row) and membrane (bottom row) based on fluorescent (left column) and virtually stained (right column) markers. (c) Correspondence between 2D instance segmentations of the nucleus and membrane based on fluorescence or virtual staining input measured using the Dice coefficient (left) and average precision (AP, right) for ∼750 cells from the test dataset. (d) Measurements of fluorescence intensity of HIST2H2BE-mNeonGreen (left) and cell area (right) based on its respective cytoplasm mask from fluorescence and virtually stained images. For a few cells, the segmentation of membrane based on fluorescence and virtual staining do not match. These FOVs appear as outliers in panel c. This occurs due to the absence of experimental label of membrane.

**Figure 4-Supplemental Movie 1.**
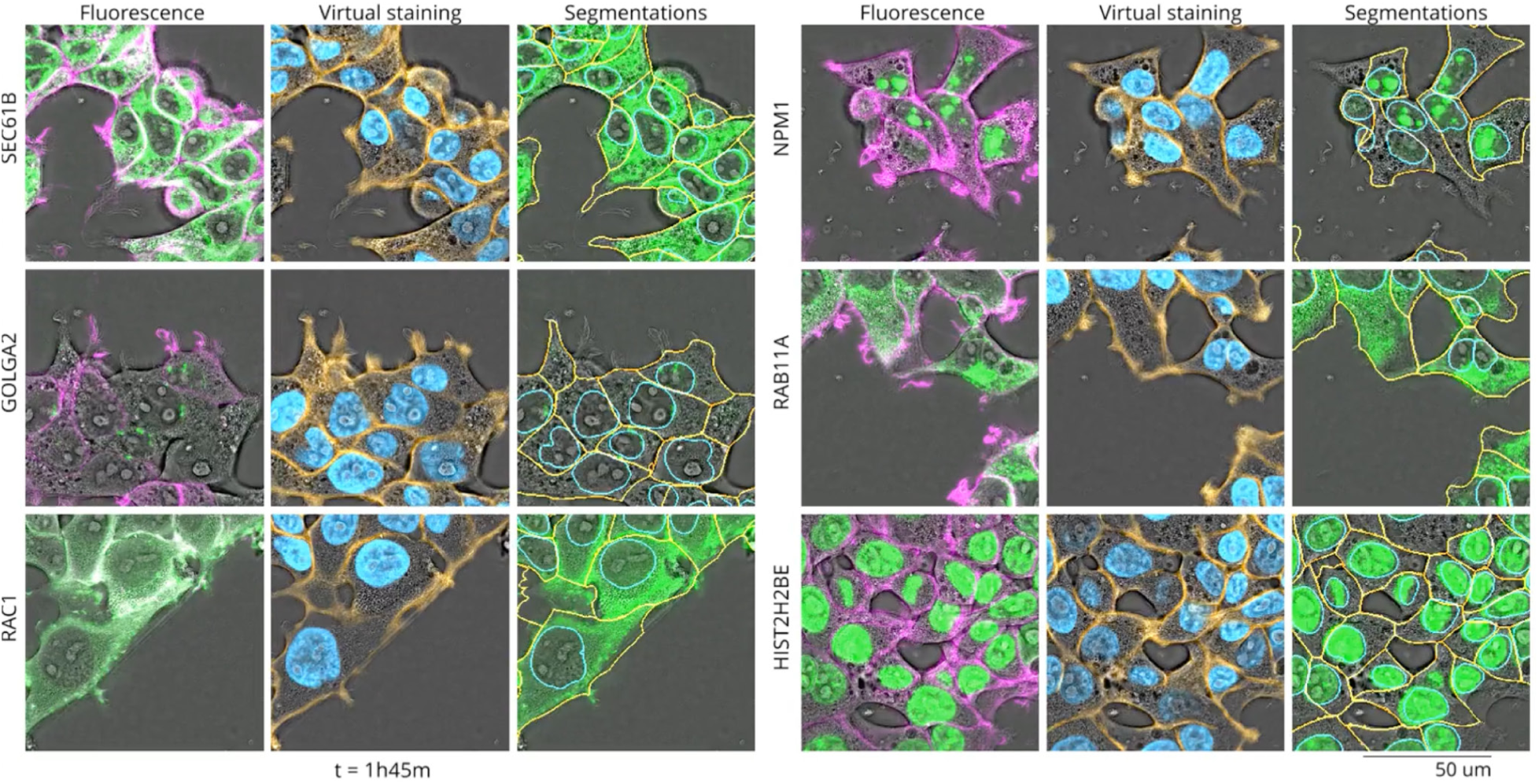
A timelapse featuring 6 OpenCell targets (SEC61B, GOLGA2, RAC1, RAB11A, NPM1, and HIST2H2BE) imaged every 15 minutes over 7 hours. Virtual staining was utilized to accurately highlight and segment landmarks such as nuclei (HIST2H2BE) and membranes (RAC1) while simultaneously visualizing other organelles (SEC61B, GOLGA2, RAB11A and NPM1) and in relation to these reference points. The targets are represented by the acquired fluorescence (membrane-magenta and organelle-green), virtual staining (membrane-yellow and nucleus-blue) an overlay of the organelles with segmentations (membrane-yellow and nucleus-blue) as contours using the in-focus slice.

**Figure 5-Supplement 1.**
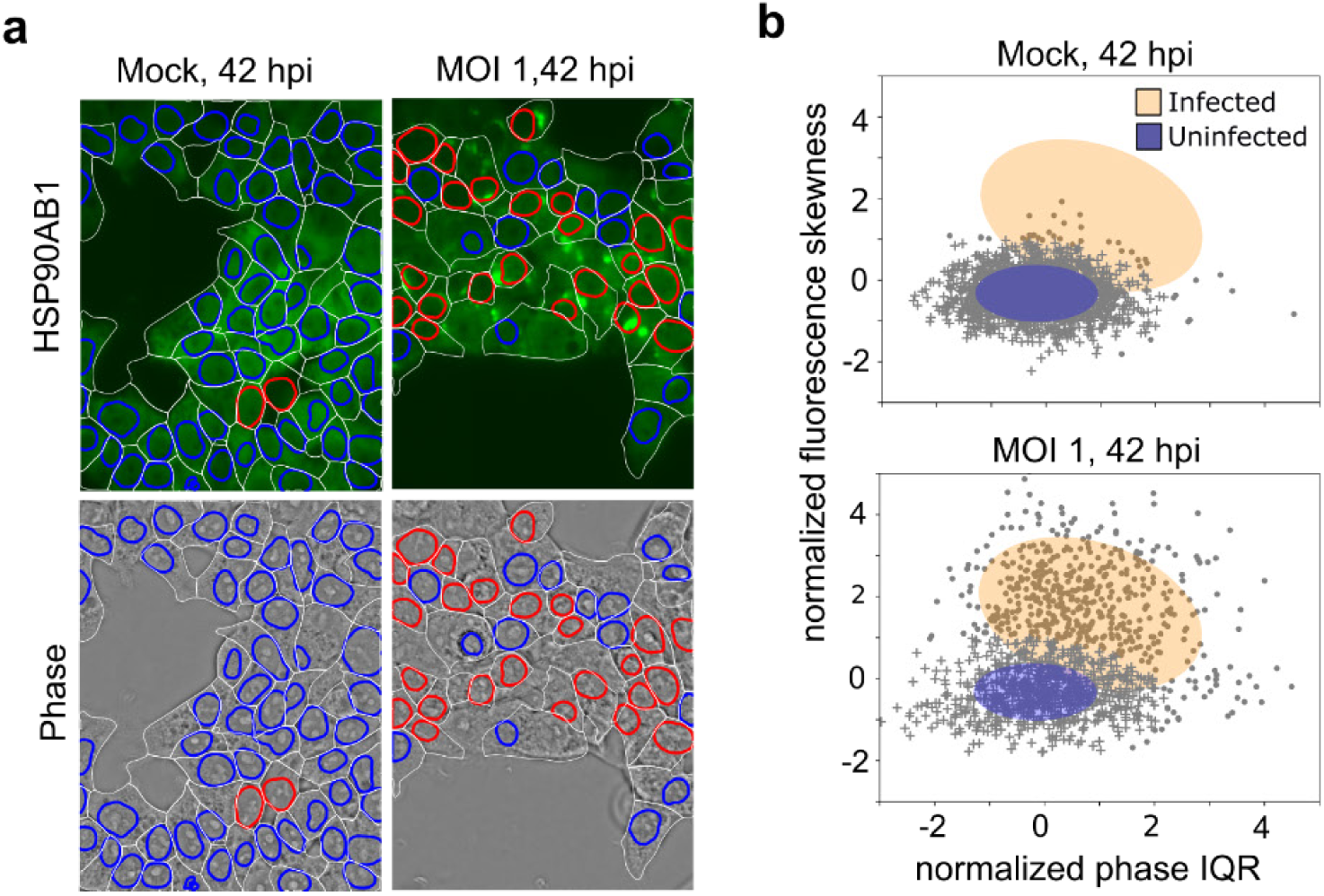
Performance of Gaussian mixture model trained on data from infected condition at 30 hpi. (a) Infection state prediction on training data used for Gaussian mixture modeling. Blue nuclear boundaries mark uninfected cells and red nuclei mark infected cells in mock (left) and MOI 1 (right) HSP90AB1 max projection (top row) and phase (bottom row) images at 42 hpi. (b) Gaussian mixture model fit on training data normalized HSP90AB and normalized phase data. Orange ellipse represents the Gaussian region containing the infected population of cells and blue region represents the uninfected cell population over mock (top) and MOI 1 (bottom) populations.

**Figure 5-Supplemental Movie 1.**
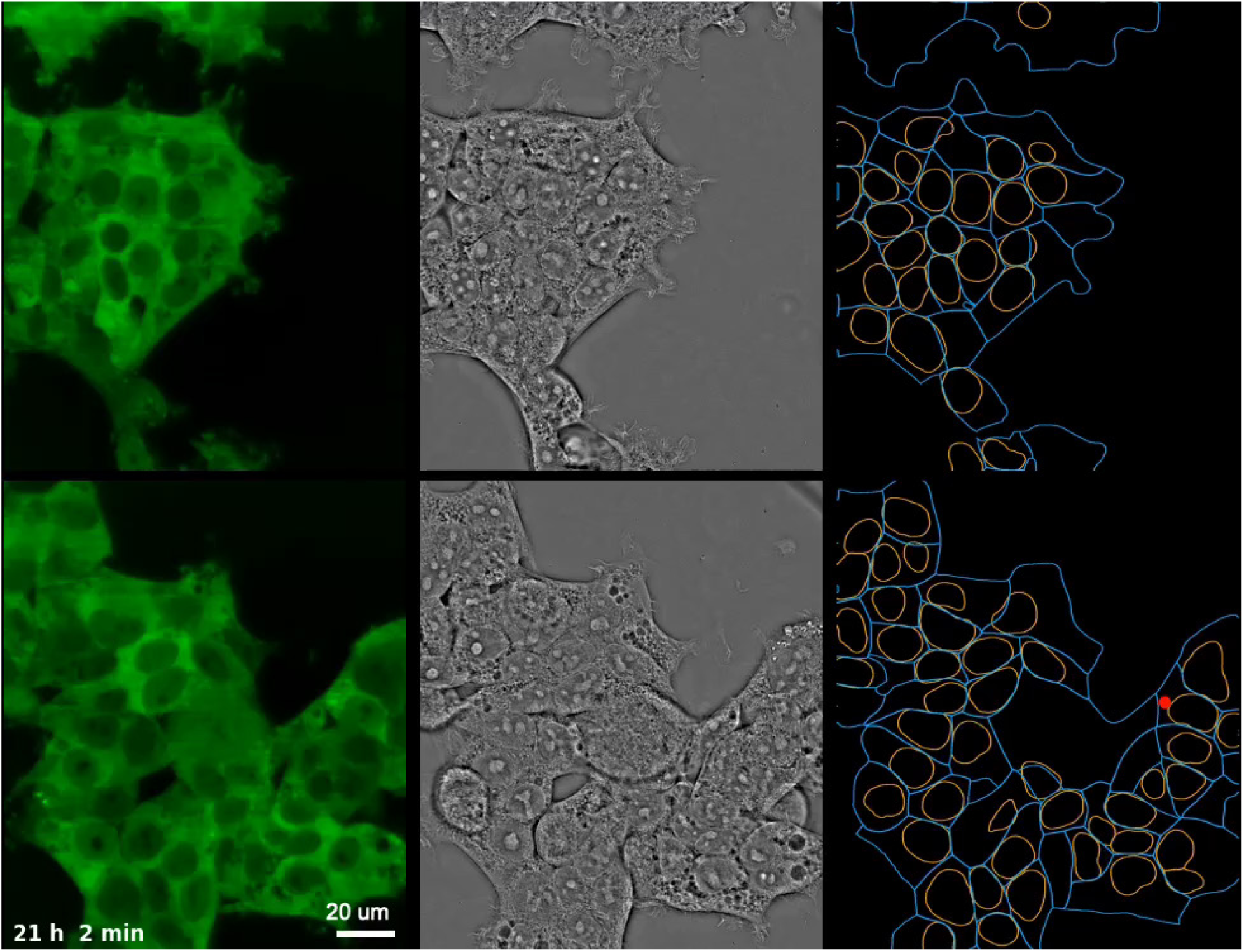
Timelapse of uninfected (top row) and infected (bottom row) HSP90AB1 cells. Shown are fluorescence (left, maximum intensity projection over 3.2 µm z-section), phase (middle), segmentation of membrane and nuclei (right, in blue and orange, respectively), and detected peaks (right, red dots).

**Figure 5-Supplemental Movie 2.**
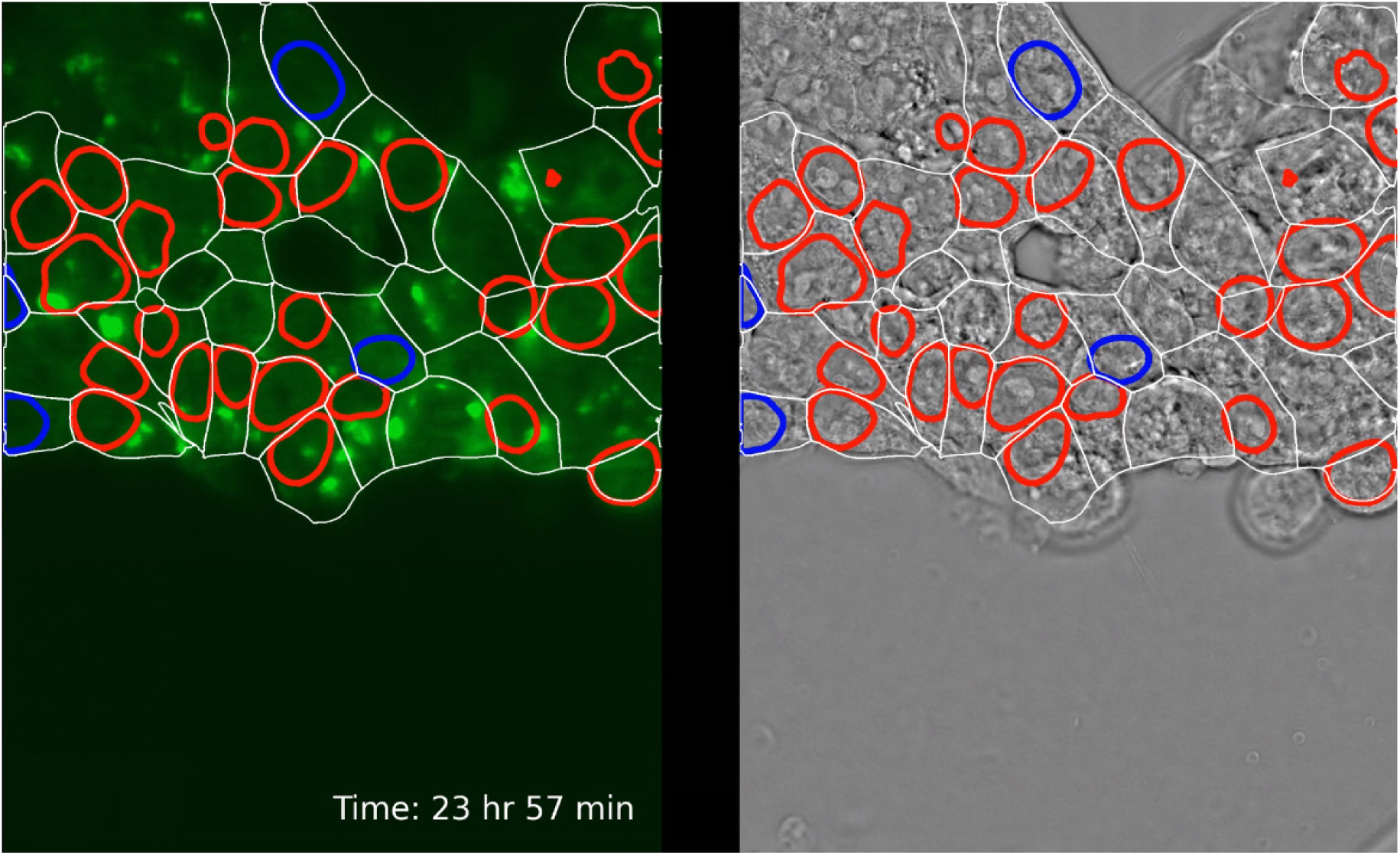
Timelapse illustrating identification of uninfected (top row) and infected (bottom row) cells. Gaussian mixture model (Figure 1-Supplement 1) was used to predict the state of infection of each cell over time. are fluorescence (left, maximum intensity projection over 3.2 µm z-section), phase (middle), segmentation of membrane (white) and nuclei of uninfected (blue) and infected (red) cells.

